# Developmental Dynamics of Naturalistic Social Behavior Revealed by Longitudinal Multi-Animal Tracking

**DOI:** 10.64898/2026.09.18.752544

**Authors:** Sunjin Kim, Yejin Jeong, Hein Lin Thant, Haeri Choi, Ain Chung

## Abstract

Early-life social experiences with parents and littermates are critical for the development of social behavior and cognition in social species. Yet, how naturalistic social interactions evolve across development remains poorly understood, because reliable tracking of multiple freely interacting animals is challenged by frequent occlusion and body overlap. To address this challenge, we developed Move-Altogether (MovAl), a multi-animal pose estimation pipeline integrating video object segmentation and contour-guided preprocessing that improved identity stability and pose accuracy compared with widely used methods. Using MovAl-derived pose sequences, we established a denoising autoencoder (DAE)-based framework to map longitudinal home-cage behavior within a common three-dimensional space defined by inter-animal spacing, social orientation, and interaction dynamics. Within this space, control mice showed progressive reorganization of group behavior from 3 to 8 weeks of age, establishing a developmental reference for naturalistic social organization. Against this reference, prenatal valproic acid (VPA) exposure produced prominent alterations in social behavior organization at 3–5 weeks of age, demonstrating that the consequences of neurodevelopmental perturbation can be detected through changes in naturalistic group behavior among littermates. Extending the analysis to preweaning mother-pup interactions revealed increased maternal movement and reduced mother-pup proximity in the VPA condition, indicating alterations in the early social environment that preceded postweaning behavioral differences. These findings suggest that quantitative characterization of early social interactions may provide a means to identify early behavioral alterations associated with atypical neurodevelopment. Together, the MovAl-DAE framework enables longitudinal quantification of naturalistic social behavior, defining its developmental organization and revealing alterations associated with neurodevelopmental perturbation.

## INTRODUCTION

Early-life social experiences with parents and littermates are critical for the normal development of social behavior and cognition (*1, 2*). In mice, social interactions begin before weaning through maternal care and interactions among littermates and continue to evolve through adolescence (*3–7*). Age-dependent changes have been observed in social investigation, social reward, and home-cage interactions with familiar cage mates (*8, 9*). Because social behavior is strongly influenced by context and partner familiarity, interactions among familiar cage mates in the home cage may capture aspects of social development that are not readily observed in conventional assays involving unfamiliar conspecifics (*6, 10–14*). Long-term group observations have revealed changes in spatial configuration, peer associations, and social hierarchies over days to weeks (*6, 15–20*). However, how social interactions within the same familiar group reorganize across development remains unclear.

A major barrier to addressing this question is reliably tracking multiple animals during naturalistic interactions. Recent multi-animal pose-estimation frameworks, including SLEAP (*21*), AlphaTracker (*22*), multi-animal DeepLabCut (*23*), and STCS (*24*) have substantially advanced keypoint detection and identity tracking through complementary strategies for pose assembly, temporal association, segmentation, and appearance- or motion-based identification. Nevertheless, these approaches can remain vulnerable during close interactions, where extensive body overlap and visual similarity obscure individual boundaries and create ambiguity in both pose assignment and identity association (*25–27*). Temporal tracking errors may also propagate across frames, while resolving fragmented or ambiguous trajectories can require post hoc reassociation or correction, challenges that become increasingly consequential in long-term recordings (*21,27,29,30*). To address these limitations, we developed Move-Altogether (MovAl), a multi-animal pose-estimation framework that preserves individual spatial information during close interactions prior to pose estimation, thereby reducing ambiguity in subsequent pose and identity assignment.

A second challenge is determining how continuous multi-animal behavior should be represented to reveal developmental organization. Existing pose-based approaches, including Keypoint-MoSeq (*28*), VAME (*31*), and B-SOiD (*32*), have been developed primarily to identify and segment recurrent behavioral motifs or states from pose dynamics, with recent approaches extending such analyses to social behavior (*33, 34*). These methods characterize behavioral repertoires and their temporal organization, while developmental comparisons would benefit from a common space capturing continuous variation in social configuration. To this end, we established a denoising autoencoder (DAE)-based framework integrating individual and inter-animal dynamics into a common behavioral space.

We applied the integrated MovAl–DAE framework to define the developmental organization of naturalistic social behavior and determine how it is altered by neurodevelopmental perturbation. Longitudinal home-cage recordings of familiar cage mates from 3 to 8 weeks of age revealed progressive reorganization from compact, low-movement configurations toward more separated and dynamic patterns. Prenatal VPA exposure, widely used to model neurodevelopmental alterations associated with ASD-like phenotypes (*35–38*), was associated with altered social behavioral organization during the early postweaning period, with group differences becoming less pronounced at later ages. Extending the analysis to preweaning mother–pup interactions further revealed increased maternal movement and reduced mother–pup proximity in the VPA condition. Together, our findings establish a longitudinal framework for characterizing how naturalistic social behavior is reorganized across development and revealing age-dependent alterations associated with prenatal neurodevelopmental perturbation.

## RESULTS

### MovAl integrates identity-preserving preprocessing with multi-animal pose estimation

To obtain stable identity-resolved pose trajectories during freely interacting multi-animal behavior, we developed Move-Altogether (MovAl), an integrated pipeline that separates identity preservation from pose estimation (Fig. 1A). Existing multi-animal pose-tracking frameworks generally infer or maintain identity through pose assembly, temporal association, or appearance-based cues during or after pose estimation (*21–24, 26, 39*). In contrast, MovAl establishes animal identity before key-point estimation by propagating individual instance masks across frames using video object segmentation. The resulting identity-preserving inputs are then used for pose estimation, reducing the need for the pose-tracking stage itself to resolve ambiguous identities during close interactions. The complete workflow comprises identity-preserving preprocessing, key-point annotation and model training, pose inference, and downstream behavioral analysis.

**Fig. 1.**
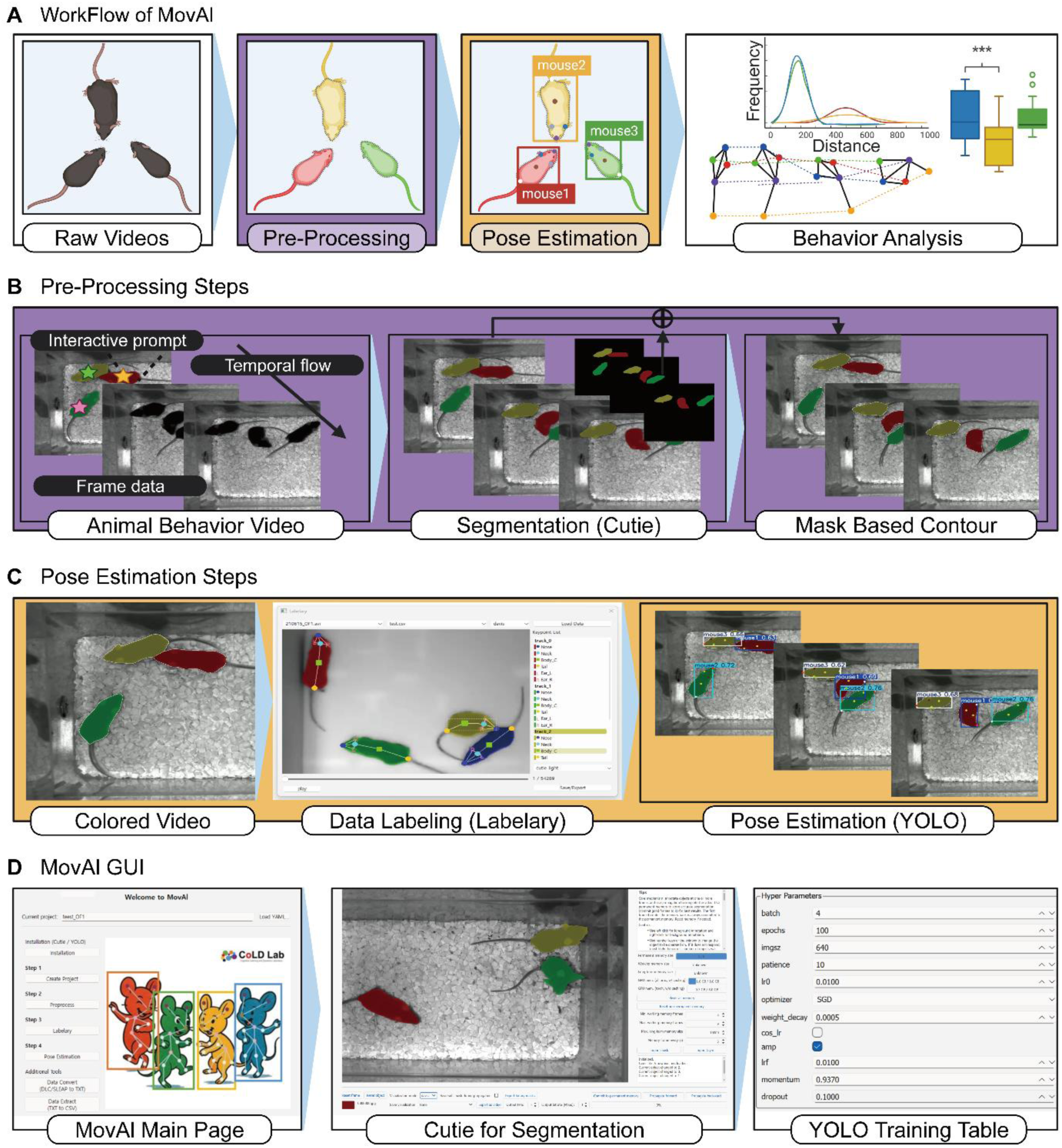
Overview of the MovAl pipeline for identity-preserving multi-animal pose estimation. **(A)** Overall workflow of MovAl. Raw multi-animal behavioral videos are processed through identity-preserving preprocessing, pose estimation, and downstream behavior analysis. **(B)** Preprocessing workflow. Raw videos are processed with Cutie, a VOS model, to propagate color-coded instance masks across frames. The propagated masks are combined with contour extraction to generate identity-preserving input videos. **(C)** Pose-estimation workflow. The identity-preserving videos are used for key-point labeling, model training, and inference. Labelary supports annotation and correction, and YOLOv11 is used as the core pose-estimation model. **(D)** MovAl graphical user interface (GUI). The GUI integrates project setup, preprocessing, Cutie-based segmentation, labeling, training configuration, and pose-estimation workflow management. Created in BioRender. Kaist, C. (2026) https://BioRender.com/g56p80r

During preprocessing, raw behavioral videos were first processed with Cutie (*40*), a video object segmentation (VOS) model that propagates initialized instance masks across successive frames (Fig. 1B). This procedure maintains a separate temporally propagated mask for each animal. The resulting instance masks were combined with contour extraction to generate identity-preserving inputs in which individual animals remained visually distinguishable throughout the recording. This preprocessing strategy was designed to reduce identity ambiguity during close social interactions, where overlap and partial occlusion make appearance-based identity assignment difficult.

The identity-preserving videos were subsequently used for key-point annotation, model training, and pose inference (Fig. 1C). YOLOv11 served as the core pose-estimation model (*41, 42*). To support annotation and iterative refinement of the pose model, we developed Labelary, a graphical labeling tool that enables keypoints to be annotated, inspected, and corrected directly on identity-preserving videos. Segmentation, annotation, model configuration, training, and inference were further integrated into the MovAl graphical user interface, providing an end-to-end workflow for processing multi-animal behavioral recordings (Fig. 1D). Together, these components were designed to generate continuous key-point trajectories in which animal identity is preserved independently of the subsequent pose-estimation step.

### MovAl improves detection continuity, identity stability, temporal stability, and accuracy

We next evaluated whether MovAl improves multi-animal pose estimation under freely interacting conditions. Five keypoints were annotated from 1,306 frames sampled across 17 top-view videos containing three mice, and SLEAP, DeepLabCut (DLC), and MovAl were trained and evaluated under raw-video and segmentation-contour input conditions (Fig. 2A). We used this benchmark to assess tracking quality across detection continuity, identity stability, temporal stability, and accuracy. Across raw-video and segmentation–contour-preprocessed conditions, SLEAP and DLC generally showed elevated tracking-miss rates across keypoints, indicating that segmentation-contour preprocessing alone did not consistently reduce tracking misses in these methods. In contrast, the full MovAl workflow maintained a mean tracking-miss rate below 1% across all annotated keypoints and significantly improved detection continuity across most comparisons with SLEAP and DLC (Fig. 2B and Table S1). For identity stability, SLEAP and DLC exhibited residual identity errors under both raw-video and segmentation–contour-preprocessed conditions, whereas the full MovAl workflow showed no identity-switch events across all 17 videos (Fig. 2C), supporting stable identity preservation in the evaluated recordings.

**Fig. 2.**
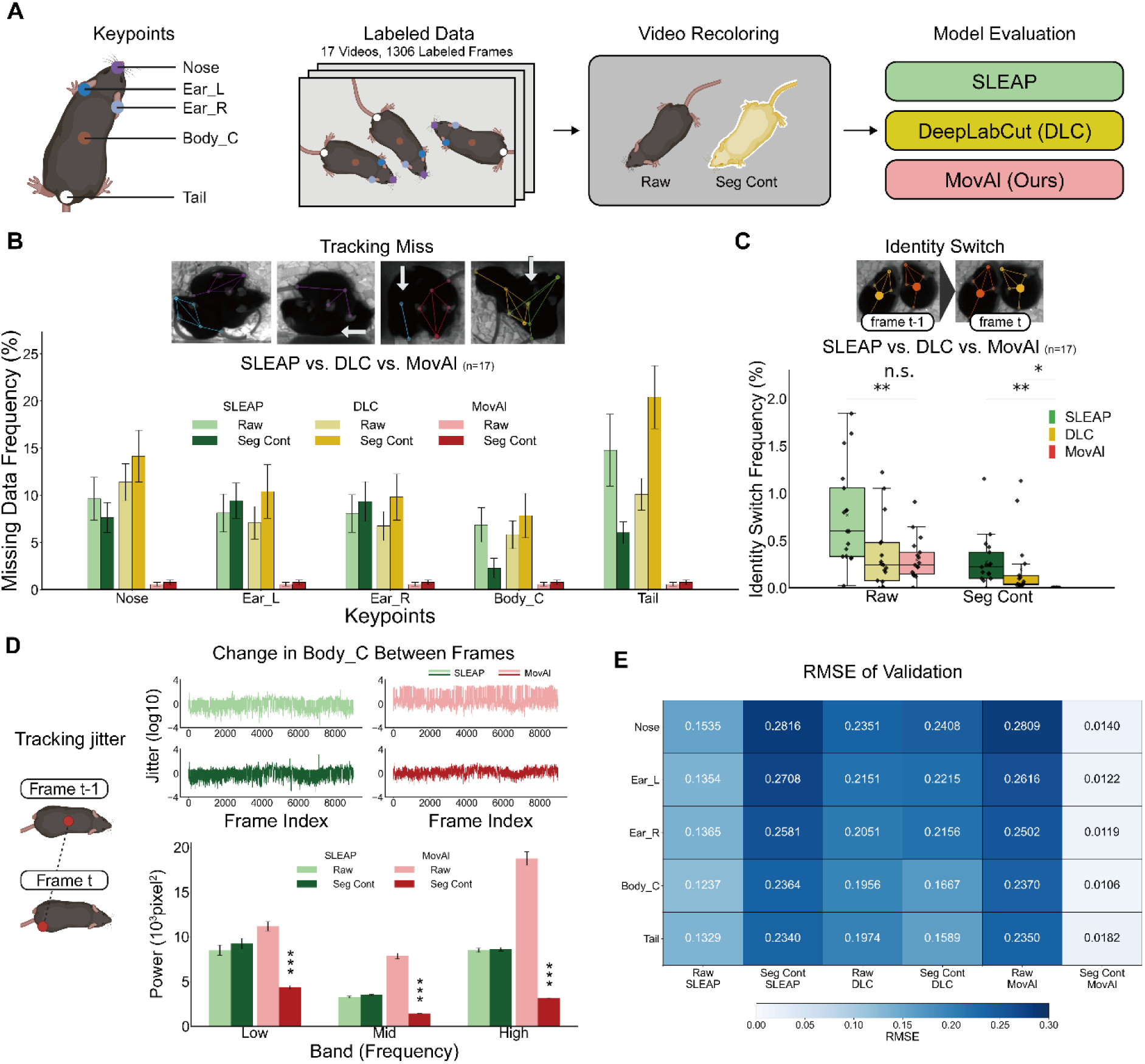
Benchmark evaluation of MovAl tracking performance. **(A)** Overview of the model-comparison experiment. Five key points were annotated: Nose, left ear (Ear_L), right ear (Ear_R), body center (Body_C), and Tail. SLEAP, DeepLabCut (DLC), and MovAl were evaluated under raw-video and segmentation-contour (Seg-Cont) input conditions. The BioRender credit applies to panel A. Created in BioRender. Kaist, C. (2026) https://BioRender.com/19belk0 **(B)** Tracking-miss frequency across annotated key points. (Top) Representative examples of missing or fragmented key-point detections. (Bottom) Percentage of frames in which each key point was not detected, shown for SLEAP, DLC, and MovAl under Raw and Seg-Cont input conditions. Bars and error bars indicate the mean ± SEM across 17 videos (two-way repeated-measures ANOVA followed by two-sided paired t-tests with Holm correction across all pairwise comparisons for each key point). Detailed Holm-adjusted *P*-values are provided in Table S1. **(C)** Identity-switch frequency. (Top) Representative images showing an identity-switch event. (Bottom) Identity-switch frequencies across methods and input conditions for 17 videos. Points represent individual videos, and box plots show the distributions (two-way repeated-measures ANOVA: input condition, *F*(1,16) = 31.65, *P* = 0.000038; method, *F*(2,32) = 14.50, *P* = 0.000033; input condition × method, *F*(2,32) = 8.83, *P* = 0.00089; two-sided paired t-tests with Benjamini–Hochberg correction for comparisons with MovAl under the corresponding input condition: Raw SLEAP, *t*(16) = 4.37, *q* = 0.0016; Raw DLC, *t*(16) = 0.64, *q* = 0.5310; Seg-Cont SLEAP, *t*(16) = 4.12, *q* = 0.0016; Seg-Cont DLC, *t*(16) = 2.32, *q* = 0.0449). \**q* < 0.05, \*\**q* < 0.01; ns, not significant. **(D)** Body-center tracking jitter. (Top) Representative log10-transformed jitter time series for each method and input condition. (Bottom) Fast Fourier transform–derived jitter power summarized across low-, mid-, and high-frequency bands. Bars and error bars indicate the mean ± SEM across 17 videos after averaging the three tracks within each video (two-way repeated-measures ANOVA: low band, method, *F*(1,16) = 7.35, *P* = 0.0154, input condition, *F*(1,16) = 29.66, *P* < 0.0001, and method × input condition, *F*(1,16) = 211.88, *P* < 0.0001; mid band, method, *F*(1,16) = 36.28, P < 0.0001, input condition, *F*(1,16) = 184.20, *P* < 0.0001, and method × input condition, *F*(1,16) = 305.59, *P* < 0.0001; high band, method, *F*(1,16) = 20.32, *P* = 0.0004, input condition, *F*(1,16) = 173.28, *P* < 0.0001, and method × input condition, *F*(1,16) = 221.03, *P* < 0.0001; two-sided paired t tests with Holm correction across the six pairwise comparisons within each band). Asterisks indicate Holm-adjusted comparisons between Seg-Cont MovAl and the other method–input combinations. Detailed Holm-adjusted P-values are provided in Table S2. \*\*\**P* < 0.001. The BioRender credit applies to the mouse schematics. Created in BioRender. Kaist, C. (2026) https://BioRender.com/19belk0 **(E)** Coordinate-level accuracy against manual annotation. Rows indicate key points, and columns indicate the method–input combinations.

To evaluate temporal stability, we calculated tracking jitter of each keypoint and compared its spectral power across predefined frequency bands. For the body-center keypoint, the full MovAl workflow reduced jitter power across low-, mid-, and high-frequency bands compared with SLEAP, with the largest reduction observed in the high-frequency band (Fig. 2D and Table S2). Similar reductions were observed for nose and tail keypoints, which represent more variable body-tip positions (Fig. S1), indicating that the improvement in tracking stability extended across multiple keypoints. Finally, to assess the spatial accuracy of each keypoint, we directly compared predicted coordinates with human annotations in the validation dataset using root mean squared error (RMSE). The full MovAl workflow showed greater agreement with manual annotations than SLEAP or DLC, with lower RMSE across all annotated keypoints, including the nose, ears, body center, and tail (Fig. 2E).

Beyond the quantitative benchmark, MovAl generated identity-resolved pose outputs for four mice in a divided chamber or an open-field arena (Fig. S2A, B and movie S1), as well as in home-cage and reflective-wall recordings (movie S1). Additional examples included six flamingos recorded from the side and two freely interacting monkeys (Fig. S2C, D and movie S2), along with five puppies (movie S2).

### Graph-temporal embedding of multi-animal home-cage behavior

After obtaining identity-resolved pose trajectories with MovAl, we developed a DAE-based model to encode each 30-frame sequence of multi-animal home-cage behavior into a 64-dimensional latent representation (Fig. 3A). Key-point coordinates were expressed relative to the group centroid to capture the animals’ relative spatial configuration, while group-centroid speed was supplied separately to retain information about collective movement.

**Fig. 3.**
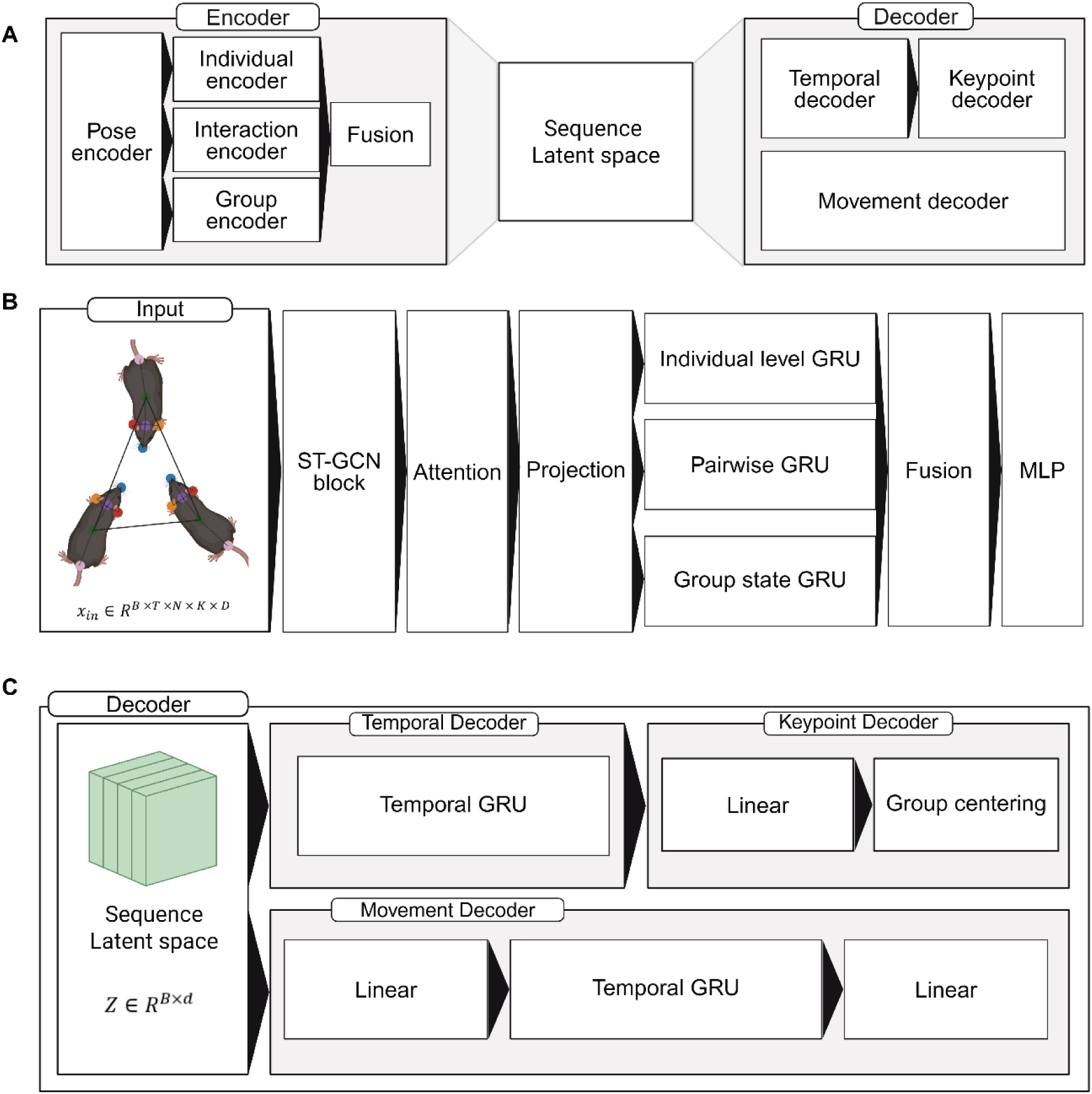
DAE-based graph-temporal representation model. **(A)** Overall architecture. Centroid-relative multi-animal pose sequences and group-centroid speed are encoded through pose, individual, pairwise-interaction, and group-state pathways. These representations are fused into a sequence-level latent embedding and decoded to reconstruct pose trajectories and group movement. **(B)** Encoder structure. A shared skeleton-based ST-GCN extracts per-animal pose features, followed by key-point attention pooling and projection. Separate bidirectional GRUs summarize individual-, pairwise-, and group-level temporal dynamics, and their outputs are fused through an MLP to produce a permutation-invariant sequence embedding. **(C)** Decoder structure. The latent representation is processed through a temporal GRU and a linear key-point decoder to reconstruct multi-animal pose trajectories, followed by group centering. A separate GRU-based movement decoder reconstructs the group-centroid speed sequence. Created in BioRender. Kaist, C. (2026) https://BioRender.com/w7s7sgn

A shared spatiotemporal graph encoder first captured anatomical relationships among connected key points and their dynamics across neighboring frames. Key-point-attention pooling then produced a pose representation for each animal at each frame. These features were used to construct individual, pairwise, and group representations. Symmetric combinations of animal features represented each pair, while pooled animal and pair features were combined with group configuration and movement features to describe the overall group state. Separate bidirectional GRUs summarized temporal dynamics at these three levels. The resulting animal and pair representations were pooled across animals and pairs, respectively, and fused with the group representation to produce the final sequence embedding, making it invariant to the arbitrary ordering of the three mice (Fig. 3B).

During training, corrupted pose inputs were encoded together with the unmodified group-centroid speed sequence. From the latent representation, a shared trajectory decoder reconstructed the three clean pose trajectories, while a separate movement decoder reconstructed group-centroid speed (Fig. 3C). This denoising objective encouraged the model to retain the underlying spatial and temporal structure of group behavior despite perturbations to the input coordinates. Training and validation losses converged over 100 epochs without a marked separation between the curves (Fig. S3A). We next tested whether the 64-dimensional bottleneck retained the relational and group-level behavioral information that the model was designed to preserve. Using nonlinear regressors evaluated by held-cage-out cross-validation, the latent representations retained information spanning spatial organization, social orientation, and movement dynamics, with high predictive performance across all examined measures (Fig. S3B). This supported the subsequent mapping of the latent representation into an interpretable behavioral space.

### Prenatal VPA exposure alters developmental trajectories of group behavior

The reliable tracking performance of MovAl enabled group-housed mice to be represented as continuous multi-animal pose sequences, which were further encoded by the DAE into sequence-level latent representations. To determine whether this framework could capture behavioral changes at the group level, we longitudinally recorded the same Control and prenatal VPA-exposed home cages from 3 to 8 weeks of age (Fig. 4A). To interpret the resulting representations, we used factor analysis of relational and group-level behavioral features from Control mice to identify three major axes (Fig. S4A). Spacing reflected inter-animal distances and spatial grouping, Orientation reflected mutual facing and approach, and Dynamics reflected relative movement and changes in group configuration (Fig. S4B). The 64-dimensional DAE representations were then mapped onto these predefined axes, providing a common latent-derived behavioral space for comparison across ages and conditions. Within this space, individual 30-frame windows of Control behavior showed a progressive age-dependent redistribution (Fig. 4B). At 3 weeks, behavioral windows were concentrated in regions of low Spacing and Dynamics, reflecting compact configurations with limited relative movement. With age, both groups occupied an expanding behavioral space characterized by greater Spacing and Dynamics and a broader range of Orientation, reflecting increased spatial separation, internal group movement, and diversity of group configurations, with VPA-exposed groups showing an earlier shift toward separated and dynamic configurations, most clearly at 4–5 weeks.

**Fig. 4.**
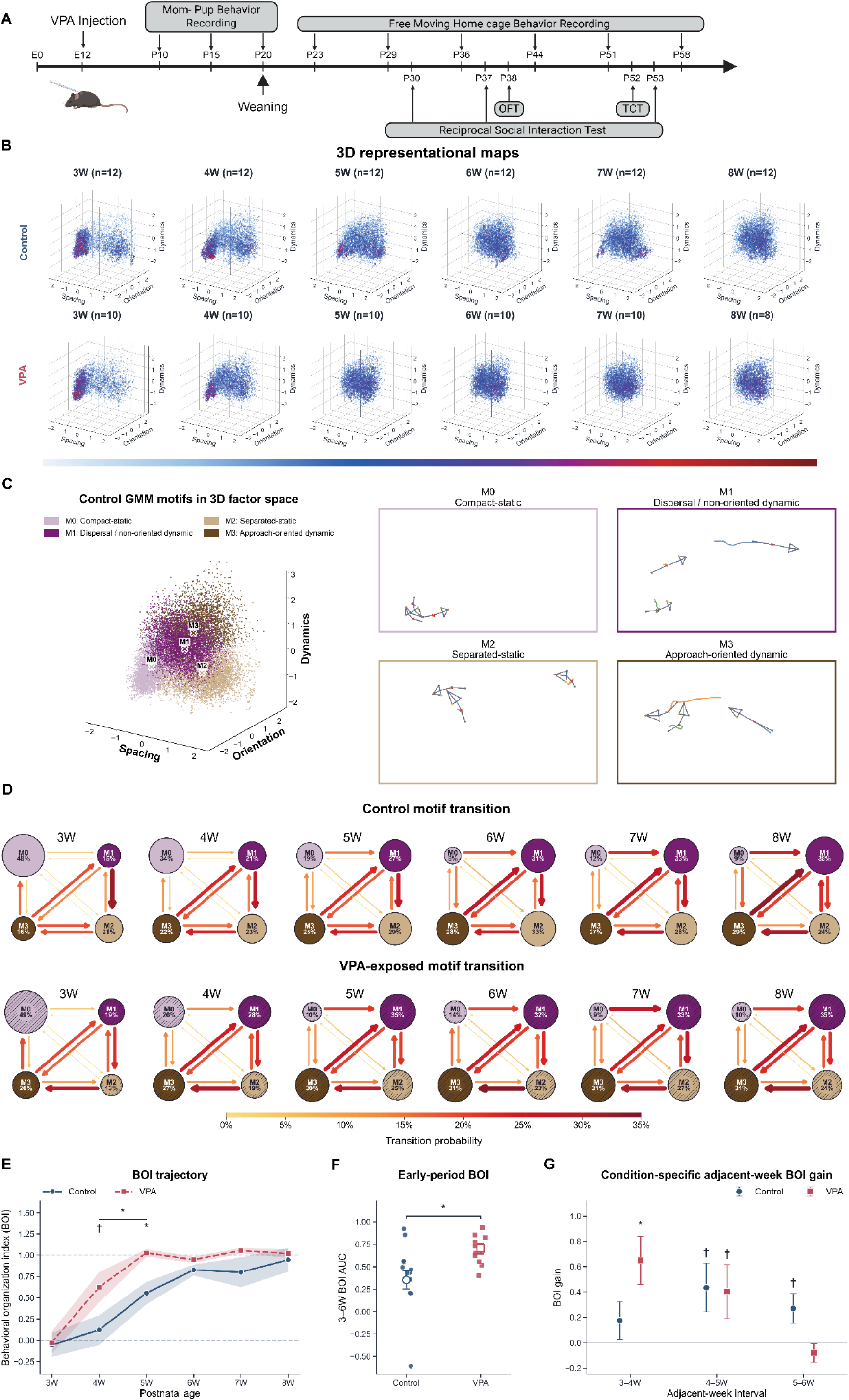
DAE-based mapping of longitudinal home-cage group behavior across adolescence. **(A)** Experimental scheme. Pregnant females in the VPA group received VPA on embryonic day 12 (E12), and mother–pup behavior was recorded on postnatal days (PNDs) 10, 15, and 20. After weaning at PND 20, freely moving home-cage behavior of the offspring was recorded longitudinally from 3 to 8 weeks of age. Reciprocal social-interaction tests were performed on PNDs 30, 37, and 53, the open-field test (OFT) on PND 38, and the three-chamber test (TCT) on PND 52. The BioRender credit applies to the schematic in panel A. Created in BioRender. Kaist, C. (2026) https://BioRender.com/w7s7sgn **(B)** Age-dependent behavioral distributions in the Control and VPA groups. Each 30-frame behavioral window was mapped onto a three-dimensional behavioral space comprising the Spacing, Orientation, and Dynamics axes. Colors from blue to red indicate increasing densities of behavioral windows (Control, *n* = 12 cages at each age; VPA, *n* = 10 cages from 3 to 7 weeks and *n* = 8 cages at 8 weeks). **(C)** Four behavioral motifs defined by applying a Gaussian mixture model (GMM) to Control behavioral windows. (Left) Each point is colored according to the motif, and X symbols indicate the center of each motif. Based on their characteristics within the behavioral space, the motifs were classified as M0, compact-static; M1, dispersal/non-oriented dynamic; M2, separated-static; and M3, approach-oriented dynamic. (Right) A representative 30-frame behavioral window for each motif. The colored lines indicate the body-center trajectories of the three mice, and the skeletons show their poses in the final frame. **(D)** Age-dependent motif transition diagram in the Control and VPA groups. Circle size and the percentage indicate the mean occupancy of each motif. Arrow direction indicates transitions between consecutive 30-frame behavioral windows, while arrow thickness and color are proportional to the conditional transition probability. The joint motif organization, incorporating both occupancy and transition probabilities, changed significantly with age in both groups (within-condition multivariate permutation tests, Holm-adjusted q < .001 for both groups). **(E)** Behavioral organization index (BOI), scaled to mean Control scores of 0 at 3 weeks and 1 at 8 weeks. Lines and shaded areas indicate estimated mean BOI ± SE from a longitudinal mixed-effects model with dam/litter/cage random intercepts and condition- and week-dependent residual variances (two-sided t-tests with Benjamini–Hochberg correction across the two weekly comparisons; VPA − Control, 4 weeks: *t*(6.01) = 2.02, *q* = 0.0894; 5 weeks: *t*(6.45) = 3.41, *q* = 0.0257). The horizontal line indicates the Control–VPA difference in mean BOI averaged over weeks 4 and 5 (two-sided t-test; VPA − Control, *t*(6.37) = 3.45, *P* = 0.0123). Standard errors and tests in E–G used dam-clustered CR1 covariance estimates and Satterthwaite degrees of freedom. \**q* < 0.05, †0.05 ≤ *q* < 0.10, \**P* < 0.05. **(F)** Normalized W3–W6 BOI AUC. Filled points represent observed cage-level AUC and open symbols show model-estimated means ± SE (Control, 0.354; VPA, 0.704; two-sided t-test; *t*(6.73) = 2.93, *P* = 0.0230). \**P* < 0.05. **(G)** Adjacent-week changes in BOI from 3 to 6 weeks, shown as model-estimated gain ± SE (one-sided t-tests against zero with Benjamini–Hochberg correction across six comparisons; 3 to 4 weeks, Control, *t*(3.50) = 1.18, *q* = 0.1877, and VPA, *t*(6.40) = 3.41, *q* = 0.0390; 4 to 5 weeks, Control, *t*(3.50) = 2.25, *q* = 0.0776, and VPA, *t*(6.40) = 1.90, *q* = 0.0776; 5 to 6 weeks, Control, 0.270 ± 0.118, *t*(3.50) = 2.30, *q* = 0.0776, and VPA, *t*(6.40) = −1.07, *q* = 0.8376). \**q* < 0.05, †0.05 ≤ *q* < 0.10.

To identify the behavioral patterns underlying this redistribution, we fitted a Gaussian mixture model (GMM) to Control behavioral windows and selected a four-motif solution that captured additional behavioral structure while maintaining high codebook stability (Fig. S5A). The four motifs occupied distinct regions of the common behavioral space (Fig. 4C), and their occupancy was calculated using soft posterior assignments, which were less sensitive than discrete labels to small shifts in behavioral-window boundaries (Fig. S5B). M0 was characterized by low Spacing and Dynamics, corresponding to a compact, relatively static configuration. M2 also showed low Dynamics but greater Spacing, representing a separated-static configuration. M1 was characterized by dispersed movement with little consistent mutual orientation, whereas M3 combined movement with increased orientation and approach toward other group members (movie S3).

The prevalence and temporal organization of these motifs changed progressively across development (Fig. 4D). At 3 weeks, the compact-static motif M0 accounted for nearly half of behavioral windows in both groups but subsequently declined in occupancy and recurrence at the group-mean level, accompanied by redistribution across M1–M3 and increased transitions from M0 to non-M0 states (Fig. S6). In Control mice, the W3–W4 interval was characterized primarily by reduced recurrence of M0 and M2 and selective changes in routing among non-M0 states, whereas reductions in M0 recurrence and increased exits from M0 became more prominent across W4–W6 (Fig. S6A). VPA-exposed groups showed a broadly similar developmental pattern, although the group means suggested that reduced M0 occupancy and recurrence and increased routing away from M0 were most apparent across W3–W5, with comparatively smaller changes across W5–W6 and apparent convergence between the two conditions in mean motif composition and major transition pathways from 6 weeks onward (Fig. S6B).

To quantify these developmental differences in group behavior, we constructed a Control-referenced behavioral organization index (BOI) from the distribution of behavioral embedding space (Fig. 4E–G). We then compared BOI trajectories between the Control and VPA groups to examine how BOI changed with age. Model-estimated mean BOI averaged over weeks 4 and 5 was significantly higher in the VPA group, indicating an overall elevation across this early postweaning interval. Separate weekly comparisons showed a trend toward higher BOI at 4 weeks and a significant difference at 5 weeks (Fig. 4E). Consistent with this elevation, normalized BOI AUC from 3 to 6 weeks was significantly greater in VPA-exposed cages than in Controls (Fig. 4F). We next examined adjacent-week BOI gains to characterize the timing of these changes. VPA-exposed cages showed a significant increase in BOI from 3 to 4 weeks and a trend toward an increase from 4 to 5 weeks. In Controls, trends toward increased BOI occurred over the successive 4–5- and 5–6-week intervals (Fig. 4G). Together, these findings indicate that VPA-exposed groups showed higher early postweaning BOI and occupied more separated and dynamic configurations, most clearly at 4–5 weeks.

Finally, to examine whether the VPA-associated elevation in BOI differed by sex, we assessed condition, sex, and condition-by-sex effects on mean BOI across 4–5 weeks and normalized BOI AUC from 3 to 6 weeks. Both measures showed a significant condition effect, with higher BOI in VPA-exposed cages when averaged across sexes. Neither measure showed a significant sex main effect or condition-by-sex interaction after multiple-comparison correction (Fig. S7). Thus, VPA-associated BOI elevations were detected, but there was no statistically supported sex effect on these measures or evidence that the VPA effect differed between males and females.

### Prenatal VPA exposure alters direct social engagement and mother–pup interactions

Following the age-dependent differences observed in home-cage group behavior, we assessed the same cohort using task-based assays to examine social behavior in other interaction contexts (Fig. 5A–D). In the three-chamber test at PND 52, both groups spent significantly more time investigating and made more visits to the novel mouse than to the familiar mouse, with preference indices significantly above zero but no significant difference between groups (Fig. 5A–C). Similarly, during the sociability phase, both Control and VPA-exposed mice spent more time investigating and made more visits to the social stimulus than to the empty cup, with significantly positive sociability preference indices in both groups (Fig. S8A, B). Additional analyses showed no significant between-group differences in total movement distance during habituation or the social-novelty phase (Fig. S8C). Neither group showed a significant left–right spatial preference during habituation, as assessed by compartment occupancy and the right-side preference index (Fig. S8D).

**Fig. 5.**
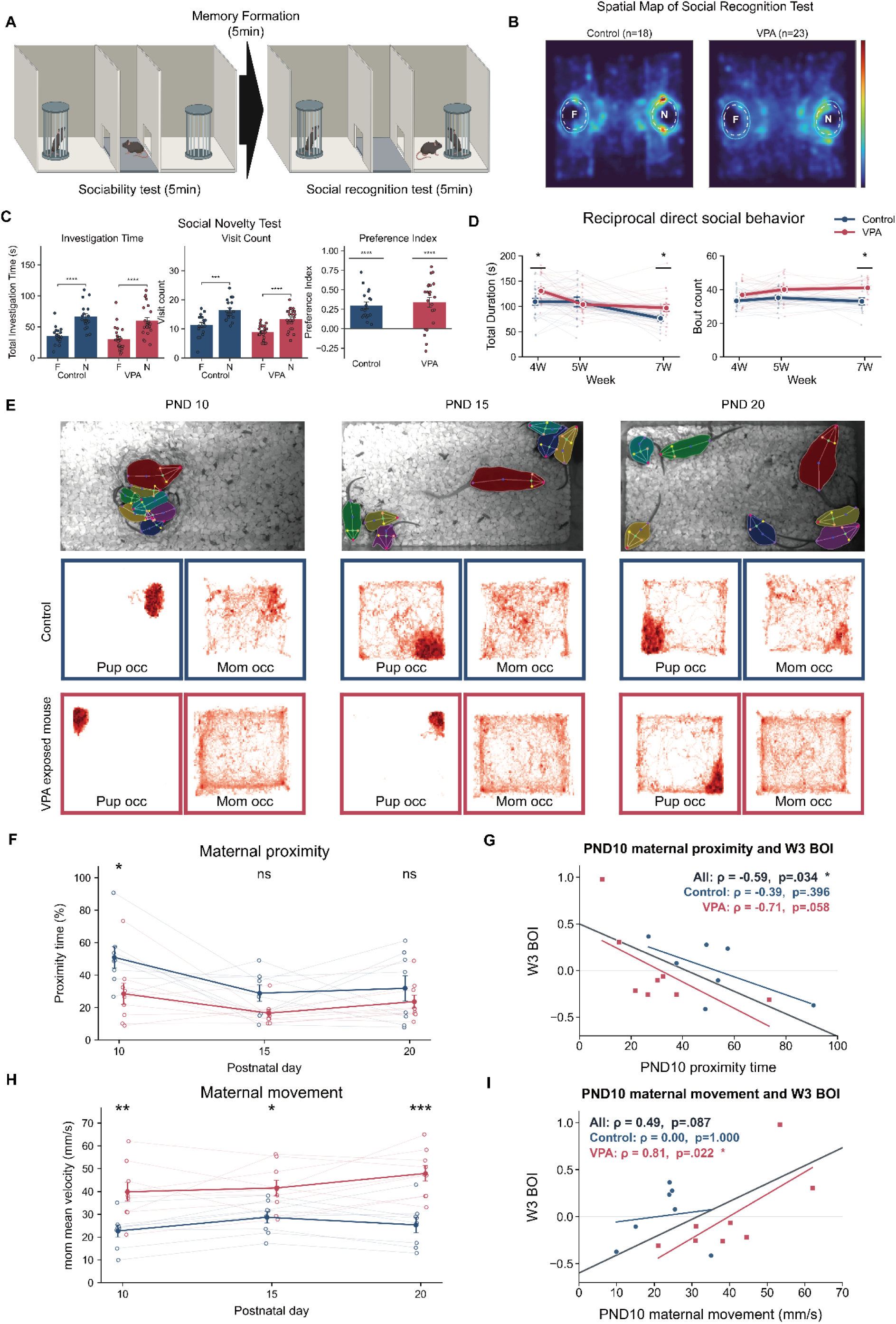
Social behavior and mother–pup interactions following prenatal VPA exposure. **(A)** Schematic of the three-chamber test. During the sociability phase, the subject investigated an empty cup and a cup containing a stranger mouse. After a 5-min interval, the familiarized mouse and a novel mouse were presented during the social-novelty phase. The BioRender credit applies to the schematic in panel A. Created in BioRender. Kaist, C. (2026) https://BioRender.com/o5j4mcj **(B)** Group-averaged spatial density maps of nose positions during the social-novelty phase. Dashed circles indicate the familiar-(F) and novel-mouse (N) regions, and colors from blue to red indicate nose-position density. **(C)** Social-novelty preference quantified by investigation time, visit count, and preference index. Each point represents one mouse, and bars and error bars indicate the mean ± SEM (Control, *n*=18; VPA, *n*=23). (Left) Total investigation time in the familiar- and novel-mouse regions (two-sided paired t-tests: Control, *t*(17)=6.84, *P*<0.0001; VPA, *t*(22)=4.82, *P*<0.0001). (Middle) Number of visits to the familiar- and novel-mouse regions (two-sided paired t-tests: Control, *t*(17)=4.91, *P*=0.0001; VPA, *t*(22)=5.10, *P*<0.0001). (Right) Preference index, calculated as ((N-F)/(N+F)), where (F) and (N) indicate the time spent in the familiar- and novel-mouse regions, respectively (two-sided one-sample t-tests against zero: Control, *t*(17)=6.83, *P*<0.0001; VPA, *t*(22)=5.43, *P*<0.0001; between-group comparison, two-sided Welch’s t-test, *t*(37.28)=−0.53, *P*=0.601). ***P<0.001 and ****P<0.0001. **(D)** Direct social behavior toward a freely moving novel conspecific at 4, 5, and 7 weeks of age. Light lines represent individual mice, and dark lines and error bars indicate the mean ± SEM (Control/VPA: 4 weeks, *n*=24/23; 5 weeks, *n*=18/21; 7 weeks, *n*=15/23). (Left) Total duration of direct social behavior (two-sided Welch’s t-tests with Benjamini– Hochberg correction across the six comparisons: 4 weeks, *t*(43.96)=−2.39, *q*=0.0475; 7 weeks, *t*(35.94)=−2.36, *q*=0.0475). (Right) Number of direct social-behavior bouts (two-sided Welch’s t-tests with Benjamini–Hochberg correction across the six comparisons: 7 weeks, *t*(20.70)=−3.58, *q*=0.0109). *q<0.05. **(E)** Application of MovAl to preweaning mother–pup recordings. Representative detections show simultaneous tracking of one mother and multiple pups at PNDs 10, 15, and 20 (top). Corresponding spatial occupancy maps are shown for pups and mothers (bottom). The upper and lower occupancy-map rows represent the Control and VPA groups, respectively, and darker colors indicate higher occupancy. **(F)** Percentage of time that the mother remained within the defined proximity threshold of at least one pup. Light lines represent individual cages, and dark lines and error bars indicate the mean ± SEM (Control, *n* = 8 cages; VPA, *n* = 9 cages; two-way repeated-measures ANOVA: condition, *F*(1,15) = 8.71, *P* = 0.00992; PND, *F*(2,30) = 5.07, *P* = 0.0126; condition × PND, *F*(2,30) = 0.92, P = 0.410; Holm-adjusted post hoc comparison: PND 10, *t*(30) = 2.84, adjusted *P* = 0.0240). *P < 0.05; ns, not significant. **(G)** Association between PND 10 maternal proximity and the BOI at 3 weeks. Each point represents one maternal litter, with BOI averaged across sibling postweaning cages where applicable. Lines indicate associations across all litters and within each condition (Control, *n* = 7; VPA, *n* = 8; All: condition-adjusted rank correlation with a two-sided exact within-condition permutation test; Control and VPA: separate Spearman correlations with two-sided exact permutation tests). \**P* < 0.05. **(H)** Mean maternal movement velocity at PNDs 10, 15, and 20. Light lines represent individual cages, and dark lines and error bars indicate the mean ± SEM (Control, *n* = 8 cages; VPA, *n* = 9 cages; two-way repeated-measures ANOVA: condition, *F*(1,15) = 21.95, *P* = 0.00029; PND, *F*(2,30) = 2.40, *P* = 0.108; condition × PND, *F*(2,30) = 1.79, *P* = 0.185; Holm-adjusted post hoc comparisons: PND 10, *t*(30) = −3.59, adjusted *P* = 0.0023; PND 15, *t*(30) = −2.69, adjusted *P* = 0.0116; PND 20, *t*(30) = −4.72, adjusted *P* = 0.00015). \**P* < 0.05, \*\**P* < 0.01, and \*\*\**P* < 0.001. **(I)** Association between PND 10 maternal movement velocity and the BOI at 3 weeks. Each point represents one maternal litter, with BOI averaged across sibling postweaning cages where applicable. Lines indicate associations across all litters and within each condition (Control, *n* = 7; VPA, *n* = 8; All: condition-adjusted rank correlation with a two-sided exact within-condition permutation test; Control and VPA: separate Spearman correlations with two-sided exact permutation tests). \**P* < 0.05.

Differences emerged when direct interaction with a freely moving novel conspecific was permitted. VPA-exposed mice showed longer total interaction durations than Controls at 4 and 7 weeks of age and more interaction bouts at 7 weeks (Fig. 5D). Attentive behaviors, reflecting orientation toward, approach to, and following of the other mouse, did not differ between groups in either cumulative duration or bout frequency at any age (Fig. S9A). In contrast, VPA mice showed greater cumulative duration and frequency of prosocial behaviors, representing active physical investigation and contact, at 4 weeks, as well as greater frequency at 7 weeks (Fig. S9B). These social differences were not accompanied by detectable changes in anxiety-related behavior in the open-field test at PND 38 (Fig. S10). Together, these assays revealed increased direct physical engagement at specific ages in VPA-exposed mice, without detectable group differences in social-novelty preference or attentive behavior.

Finally, we extended the analysis to preweaning mother–pup interactions to characterize the early social environment. MovAl simultaneously tracked mothers and pups at PNDs 10, 15, and 20 despite substantial size differences and frequent overlap, enabling quantification of spatial occupancy, mother–pup proximity, and maternal movement (Fig. 5E and movie S4). At PND 10, mothers in the VPA group spent significantly less time near at least one pup than Control mothers (Fig. 5F). Across litters, this early proximity measure was negatively associated with offspring BOI at 3 weeks after accounting for condition (Fig. 5G). Mothers in the VPA condition also showed higher mean movement velocity at PNDs 10, 15, and 20 (Fig. 5H). Within the VPA group, higher maternal movement velocity at PND 10 was associated with higher offspring BOI at 3 weeks, whereas this association was not significant within the Control group or in the condition-adjusted analysis across all litters (Fig. 5I).

## DISCUSSION

In this study, we developed the MovAl–DAE framework to track naturalistic group behavior and characterize its organization across development. Control mice shifted from close, low-movement configurations after weaning toward more separated and dynamic patterns with age. Prenatal VPA exposure was associated with higher Control-referenced BOI scores during the early postweaning period, indicating an alteration in developmental behavioral organization. Alterations in mother-pup interactions were also prominent and were associated with offspring group behavior in early development. Together, these findings highlight age-dependent changes in social behavioral organization associated with prenatal neurodevelopmental perturbation.

A key technical contribution of MovAl was preserving animal identity before pose estimation. Although existing multi-animal pose-estimation methods have enabled detailed tracking, stable identity assignment remains difficult when visually similar animals cluster or occlude one another, even with complementary strategies such as temporal association, segmentation, motion cues, and appearance-based re-identification (*21, 23, 24, 43–45*). MovAl takes a different approach by propagating identity-specific object masks through time and using these masks to generate identity-constrained inputs for pose estimation. This separation of identity preservation from pose estimation improved identity continuity and pose stability during close interactions, reducing tracking artifacts that could otherwise distort estimates of inter-animal relationships.

Beyond tracking, pose-based representation methods have enabled complex behavior to be organized into recurrent motifs or states and have revealed behavioral differences across experimental conditions (*24, 28, 31, 32*). Such approaches are particularly useful for identifying discrete or condition-associated behavioral patterns, whereas our aim was to characterize how the organization of spontaneous social behavior changes continuously across development. We therefore constructed a common behavioral space that remained fixed across ages and conditions, enabling developmental reorganization to be quantified longitudinally and providing a common reference for evaluating age-dependent effects of neurodevelopmental perturbation.

Using this framework, we observed a postweaning shift in Control groups from predominantly compact-static behavior at 3 weeks toward separated-static and dynamic states with age. This shift involved changes in both movement dynamics and the spatial organization of static behavior. Previous home-cage studies likewise reported high levels of huddling in juvenile C57BL/6J mice, followed by declining huddling and increasing sociopositive interactions toward adulthood (*6, 46, 47*). Within this broader developmental shift, Control BOI showed a trend toward an increase between 4 and 5 weeks, corresponding to PNDs 29–36 and spanning the peri-pubertal period in C57BL/6J mice (*48, 49*). This interval has also been associated with experience-dependent maturation of prefrontal circuits and with heightened sensitivity to post-weaning peer experience (*50–53*). Because huddling contributes to thermoregulation and also provides sustained close social contact (*54–57*), this transition may reflect a shift from an early organization characterized by frequent physical proximity toward one combining spatially independent rest with a broader range of movement and approach states.

VPA-associated BOI elevations were most apparent during the early postweaning period, with mean values becoming more similar between groups from 6 weeks onward. Beyond this age dependence, the behavioral differences observed following VPA exposure also varied across interaction contexts. VPA-exposed mice retained social-novelty preference in the three-chamber test at PND 52. The preserved social-novelty preference is consistent with some previous studies of adult mice prenatally exposed to VPA (*58–60*), although others have reported reduced or absent preference (*61–64*). In the direct-interaction assay, no significant group differences were detected in attentive behaviors, including approach, facing, and following, whereas behaviors categorized as prosocial, including physical investigation and contact, increased at specific ages. This pattern suggests that VPA-associated differences were more apparent in direct physical engagement than in the measured orienting and approach behaviors.

These findings underscore the complementary information provided by different social assays. The three-chamber test assesses preference between spatially constrained stimuli and limits direct physical interaction with the stimulus mouse (*12, 65*), potentially limiting its ability to capture the differences observed during freely moving reciprocal interactions. Longitudinal home-cage monitoring enables behavioral changes to be examined across development (*66, 67*). Although home-cage organization and social-novelty preference represent distinct aspects of social behavior, our findings illustrate how longitudinal analysis can complement assessment at a single later age by identifying developmental periods during which behavioral alterations are most apparent.

Differences were also evident before weaning, with increased maternal movement and reduced mother–pup proximity in the VPA condition. Mother–pup interactions are bidirectional, and prenatally VPA-exposed pups have been reported to show altered ultrasonic vocalizations and responses to maternal or familiar olfactory cues (*68–71*). Therefore, our results may reflect alterations in maternal behavior, pup signaling, or their reciprocal relationships. PND 10 maternal proximity was associated with offspring BOI at 3 weeks, while maternal movement showed an additional association within the VPA group.

Our findings provide a framework for defining developmental changes in social organization while also opening several questions about their underlying mechanisms. Although early mother– pup interactions were associated with subsequent behavioral organization, the present study does not establish a causal relationship between early social experience and later group behavior. The BOI should be interpreted as a relative measure of behavioral organization rather than an absolute index of developmental maturity, and the behavioral states identified here describe patterns of movement and social configuration without assigning specific behavioral functions or motivations. Technically, the sequential segmentation and pose-estimation pipeline and offline behavioral analysis currently limit real-time applications. Despite these limitations, the MovAl– DAE framework provides a basis for investigating how early social interactions relate to subsequent developmental changes in group behavior.

## MATERIALS AND METHODS

### Animals and prenatal VPA exposure

Control and prenatally VPA-exposed C57BL/6J mice were used in this study. Female breeders aged 8–24 weeks were mated under specific-pathogen-free conditions, and vaginal plugs were checked daily. The day on which a vaginal plug was detected was designated embryonic day 0 (E0). At E12, pregnant females in the VPA group received a subcutaneous injection of valproic acid sodium salt dissolved in 0.9% saline at a dose of 600 mg/kg. The solution was prepared at 50 mg/mL and adjusted to pH 7.3, with the injection volume calculated according to body weight. Pregnant females in the Control group received no treatment. All animal procedures were approved by the Institutional Animal Care and Use Committee of the Korea Advanced Institute of Science and Technology (KAIST; approval no. KA2025-006-v7).

### Longitudinal home-cage behavior recording

Home-cage behavior was recorded longitudinally before and after weaning. Mother–pup interactions were recorded at postnatal days (PNDs) 10, 15, and 20. Following weaning at PND 20, recordings continued with three same-sex offspring housed together per cage from 3 to 8 weeks of age. For each session, animals were transferred from their housing cage to a separate MVCS recording cage (200 × 320 × 145 mm, width × depth × height) containing white square pulp-chip bedding to improve contrast between the animals and the background. Animals were habituated for at least 1 h in the recording cage within a sound-attenuated behavioral chamber. The standard wire-grid lid was replaced with a flat recording cover to prevent escape during habituation and recording. All sessions were conducted during the dark phase after 8 p.m., with the lights off and infrared illumination. Following habituation, freely moving behavior was recorded for 15 min using an overhead infrared-sensitive machine-vision camera (MU-A121R-31, Crevis) at 30 frames/s and a resolution of 1920 × 1080 pixels.

### Mother-pup interaction

Home-cage videos recorded at PNDs 10, 15, and 20 were analyzed using MovAl to track the mother and individual pups. Normalized coordinates were converted to millimeters using the cage dimensions (320 × 200 mm). Frames were considered valid for proximity analysis when the mother and at least one pup had valid body-center coordinates with confidence scores ≥0.5. Mother–pup distance was calculated in each valid frame as the distance between the mother’s body center and that of the nearest detected pup. For each recording, the proximity threshold was defined as the mean of the mother’s median nose–tail length and the median nose–tail length pooled across all valid pup measurements. A proximity bout required this distance to remain within the threshold for at least 1 s, regardless of the identity of the nearest pup. Any invalid frame terminated the bout; missing intervals were neither bridged nor interpolated. Maternal proximity was expressed as the percentage of valid tracking time occupied by qualifying bouts. Mean maternal movement velocity was calculated as the total body-center displacement across consecutive valid frames divided by the corresponding tracking time. Frame-to-frame displacements exceeding 50 mm were excluded as tracking errors.

### Direct social-interaction test

The direct social-interaction test assessed social engagement toward a freely moving novel conspecific at PNDs 30, 37, and 53. Before testing, each test mouse was individually habituated for 1 h in an MVCS cage containing white pulp-chip bedding within a sound-attenuated behavioral chamber. Stimulus mice were habituated separately for 1 h and were unfamiliar C57BL/6J wild-type mice younger than the corresponding test mouse by no more than 1 week. Following habituation, one stimulus mouse was introduced into the test mouse’s cage, and social interaction was recorded for 5 min. Recordings were performed in the dark using infrared illumination and an infrared-sensitive machine-vision camera. Only behaviors directed by the test mouse toward the stimulus mouse were quantified.

Videos were analyzed using MovAl-derived pose outputs and random forest classifiers implemented in SimBA. Behavioral bouts were manually annotated in a subset of videos by marking the onset and offset of each behavior, and these annotations were used to train the classifiers. The trained classifiers were then applied to the remaining videos using a classifier probability threshold of 0.5. Overall social behavior comprised Approach, Facing, Following, Nose–Head, Nose–Body, Nose–Anogenital, and Mounting. Approach, Facing, and Following were grouped as attentive behavior, whereas Nose–Head, Nose–Body, Nose–Anogenital, and Mounting were grouped as prosocial behavior. For each category, frames were classified as positive when at least one constituent behavior was detected. Positive segments separated by interruptions shorter than 15 frames were merged, and merged bouts lasting at least 15 frames were retained. Cumulative duration and bout count were calculated as the total duration and number of retained bouts, respectively, for each mouse in each recording.

### Three-chamber test

The three-chamber test was performed at PND 52 to assess sociability and social-novelty preference under illumination maintained at 30 lux. Stimulus mice were unfamiliar C57BL/6J wild-type mice aged 7–11 weeks and were of the same sex as the test mouse. Before testing, test and stimulus mice were habituated for 1 h in separate compartments of a sound-attenuated behavioral chamber. During apparatus habituation, each test mouse was confined to the center chamber for 5 min and then allowed to explore all three chambers for 5 min with both wire cups empty. Cylindrical wire cups measured 90 mm in internal diameter and 140 mm in height.

During the sociability phase, a novel stimulus mouse was placed in one wire cup, while the opposite cup remained empty. The test mouse was allowed to explore the apparatus for 10 min. After a 5-min resting period in the center chamber, the social-novelty phase began with a second novel stimulus mouse placed in the previously empty cup and the first stimulus mouse remaining as the familiar stimulus. The test mouse was again allowed to explore for 10 min. The chamber floor and cups were cleaned with ethanol after testing each mouse.

Quantitative analyses used the first 5 min of each test phase. Each cup boundary was manually outlined as a polygon, which was expanded outward by 20% of the cup’s mean radius to define the investigation region of interest (ROI). Frames in which the test mouse’s nose was within the ROI were classified as investigation-positive. Interruptions of 15 frames or fewer were bridged within each cup-specific investigation sequence, after which positive bouts shorter than 15 frames were discarded. At 30 frames/s, these thresholds corresponded to 0.5 s. Investigation time was calculated as the total number of positive frames after processing divided by the frame rate. Each retained positive bout was counted as one visit, including bouts beginning at the start of the analysis period. The sociability preference index was calculated as (T_S_ – T_E_) / (T_S_ + T_E_), where T_S_ and T_E_ denote investigation times for the social and empty cups, respectively. The social-novelty preference index was calculated as (T_N_ – T_F_) / (T_N_ + T_F_), where T_N_ and T_F_ denote investigation times for the novel and familiar mice, respectively. Individual mice were treated as the experimental units.

### Open field test

The open-field test was performed at PND 38. Before testing, each mouse was habituated individually for 1 h in a sound-attenuated behavioral recording chamber. Each mouse was then placed in a 40 × 40 cm white acrylic arena, and freely moving behavior was recorded for 10 min.

MovAl-derived body-center coordinates were used to quantify movement and spatial occupancy. The center zone was defined as the central 20 × 20 cm region of the arena. Total distance traveled, distance traveled within the center zone, and time spent in the center zone were calculated. Normalized coordinates were converted to centimeters using the arena dimensions, and frame-to-frame displacements exceeding 5 cm were excluded as tracking errors. Individual mice were treated as the experimental units.

### MovAl pose-estimation pipeline

Raw behavioral videos were processed using the Move-Altogether (MovAl) pipeline to generate identity-resolved keypoint sequences. Individual animals were initialized using user-provided prompts and tracked across frames using the Cutie video object segmentation model (*40*). The resulting identity-specific masks and contours were used to generate inputs for pose estimation. Keypoints were annotated and corrected using Labelary, and YOLOv11 pose models trained on these annotations were applied throughout each recording (*41, 42*). The MovAl pipeline was implemented in Python 3.9 using PyTorch 2.1.2 and CUDA 12.1 and was run on an NVIDIA GeForce RTX 3060 GPU.

### Pose-estimation benchmark

To train and evaluate the pose-estimation models, 1,306 frames sampled from 17 top-view videos of three freely interacting mice were manually annotated for five key points: the nose, left ear, right ear, body center, and tail. SLEAP, DeepLabCut (DLC), and the YOLOv11-based MovAl model were trained for 200 epochs using either raw frames (Raw) or segmentation-contour-preprocessed frames (Seg-Cont) and applied to all 17 videos. Tracking-miss frequencies were averaged across the three tracks within each video and summarized as the mean ± SEM across 17 videos, with each video serving as the unit of statistical analysis.

Tracking jitter was quantified as the frame-to-frame Euclidean displacement of each keypoint in pixel coordinates. Displacements were first calculated between consecutive frames, after which values involving missing predictions and zero displacements were excluded. The remaining displacements were concatenated in their original order, log10-transformed, and mean-centered before fast Fourier transform (FFT) analysis. No interpolation was performed. Spectral power was calculated from the squared FFT magnitude and summarized within low (0.00–0.05), mid (0.05– 0.15), and high (0.15–0.50) frequency bands, expressed in cycles per retained sample. Because excluded values were removed before FFT analysis, these frequencies describe variation across retained samples rather than the original recording time.

Keypoint localization accuracy was evaluated using a held-out set of 1,250 manually annotated frames excluded from model training. Two benchmark videos were downsampled from 30 to 8 fps and divided into non-overlapping 250-frame clips, of which five were randomly selected and manually annotated, yielding 1,250 frames excluded from model training. After normalizing horizontal and vertical coordinates by image width and height, respectively, root mean squared error (RMSE) was calculated for each keypoint and method–input combination by pooling valid frame–animal pairs across all five clips. Zero coordinates were treated as missing, and pairs with missing predicted or annotated coordinates were excluded.

### Pose sequence preprocessing

MovAl-derived pose sequences contained six keypoints—the nose, body center, left ear, right ear, neck, and tail—for each of three mice. Keypoint predictions with confidence scores below 0.5 were treated as missing, and gaps of up to four consecutive frames bounded by valid coordinates on both sides were linearly interpolated. Tracking validity was assessed separately for each mouse using observed keypoints, excluding interpolated points. Tracking was marked as invalid from the start of any sequence of five or more consecutive frames in which the body center was missing or fewer than four keypoints were detected. To restore valid tracking, the neck, body center, and tail plus at least one additional keypoint had to be detected for five consecutive frames, with body length remaining within 0.5–2.0 times the mouse’s recording-level reference length. Recordings acquired at 30 frames/s were divided into non-overlapping 30-frame windows corresponding to 1 s of behavior. Each 1-s window was excluded if tracking was invalid for any mouse at any point within the window. Windows were also excluded if missing or infinite values remained in the model inputs after interpolation.

For each recording, the four cage-wall ROI corners were mapped to a unit square using a projective transformation. The x coordinates were then scaled by the video aspect ratio (1920/1080), while y coordinates were unchanged. Body length was defined as the sum of the neck-to-body-center and body-center-to-tail distances. After interpolation, median body length was calculated for each mouse across the recording. All coordinates were divided by the median of these three values. The group center was calculated in each frame as the mean body-center position of the three mice and subtracted from all keypoint coordinates.

### Pose-derived behavioral features

Pair- and group-level behavioral features were calculated at each frame from body-length-normalized pose coordinates. Let c_i,t_, n_i,t_, k_i,t_, and q_i,t_ denote the body-center, nose, neck, and tail coordinates of animal i at frame t, respectively, and let v_i,t_denote its body-center velocity. Distances were expressed in body lengths and velocities in body lengths per second. Temporal derivatives were estimated using finite differences across neighboring frames.

Pairwise proximity was represented by body-center distance, nose-to-nose distance, and the two directional nose-to-tail-base distances. These quantities were calculated as Euclidean distances and transformed using log(1 + x) to reduce the influence of large values. Directional facing from animal i toward animal j was defined as the cosine similarity between the neck-to-nose vector of animal i and the vector connecting its body center to that of animal j,

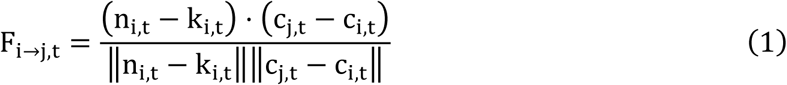

Facing was calculated in both directions for each pair, with higher values indicating stronger alignment toward the other animal. Relative movement was described by approach rate and relative speed. Approach rate was calculated as the negative projection of relative velocity onto the direction connecting the two body centers,

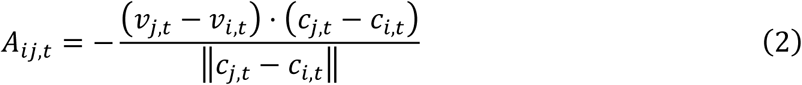

Relative speed was calculated as

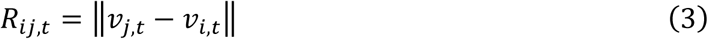

Positive and negative approach-rate values indicate approach and withdrawal, respectively. Approach rate was transformed using the inverse hyperbolic sine to preserve its sign, whereas relative speed was transformed using log(1 + x).

Group spatial configuration was derived from the three pairwise body-center distances. Overall group spacing was calculated as log(1 + mean pairwise distance). To quantify configurations in which two animals were closer to each other than to the third, dyadic grouping was defined as

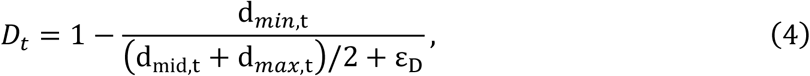

where d*_min_*_,t_, d_mid,t_, and d*_max_*_,t_ are the three pairwise distances in ascending order, and ε_D_ = 10^−6^. Values were clipped to the range 0–1. Values approaching 0 indicate similar distances among all three animals, whereas values approaching 1 indicate a relatively close pair and a more distant third animal.

Configuration speed quantified changes in the pairwise distances within the group,

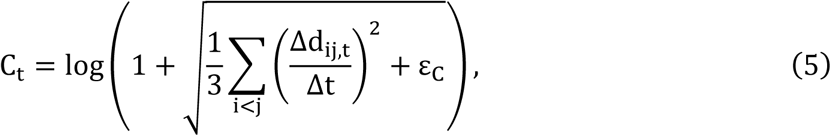

where d_ij,t_ is the body-center distance between animals i and j, and ε_C_ = 10^−8^.

Group-centroid speed was calculated from body-length-normalized coordinates before frame-wise centering,

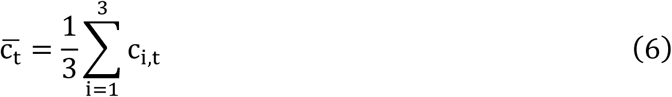

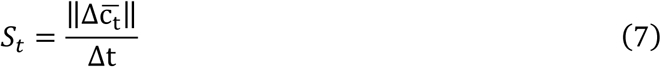

Here, c̅_t_ is the mean body-center position of the three mice. Configuration speed captures changes in inter-animal distances, whereas group-centroid speed captures movement of the group as a whole.

### Architecture of DAE-based behavioral embedding model

A denoising autoencoder (DAE) (*72*) was trained to generate one behavioral embedding per 30-frame pose window. Inputs consisted of group-centered keypoint coordinates with dimensions 30 × 3 × 6 × 2 (frames × mice × keypoints × coordinates) and a 30-frame group-center speed sequence. During training, noise was added only to the pose coordinates. The unmodified coordinates and group-center speed served as reconstruction targets.

Each mouse was represented by a six-node skeleton graph. Edges connected the nose to the neck and both ears, each ear to the neck, the neck to the body center, and the body center to the tail. A shared two-block spatiotemporal graph-convolutional encoder (*73*) processed each mouse independently, without edges between animals. Attention pooling combined the keypoint features, which were then projected into a 64-dimensional pose token for each mouse at each frame.

For each of the three animal pairs, the two pose tokens were combined with the relational features defined above. Shared weights and symmetric mean and maximum pooling were used to construct pair tokens that were invariant to animal order. A group token was formed by combining pooled individual and pair representations with group-configuration features and group-center speed.

Temporal dynamics were encoded separately at the individual, pair, and group levels using single-layer bidirectional gated recurrent units (GRUs), with 64 hidden units in each direction (*74, 75*). The individual and pair representations were each pooled across animals or pairs using element-wise mean and maximum operations. These pooled representations were combined with the group representation and passed through a multilayer perceptron (MLP) to produce a 64-dimensional latent embedding for each window. A shared GRU-based decoder reconstructed the three pose trajectories from this embedding, using a learned output-specific embedding for each trajectory. A separate GRU-based decoder reconstructed the 30-frame group-centroid speed sequence from the latent embedding using a linear output layer followed by a softplus activation to ensure nonnegative outputs.

### Training method of DAE-based behavioral embedding model

The DAE was trained using a Control-only reference dataset comprising longitudinal Control recordings and additional home-cage recordings. The dataset included 26 cages housing female mice (n = 14) or male mice (n = 12). Recordings were available from 21 cages at 3 weeks (11 female and 10 male), 21 cages at each of 4 and 5 weeks (10 female and 11 male), 19 cages at 6 weeks (9 female and 10 male), and 20 cages at each of 7 and 8 weeks (10 female and 10 male). The dataset was divided into 21 training cages and 5 validation cages, with all recordings from each cage assigned to the same subset.

The model was trained to reconstruct the clean pose sequence and group-centroid speed from 30-frame windows. To account for arbitrary animal ordering, pose reconstruction loss was calculated as the minimum mean squared error across all six possible assignments of the three reconstructed trajectories to the target trajectories. Huber loss was used for group-centroid speed reconstruction. After aligning the reconstructed trajectories with the targets, pairwise and group-configuration features were calculated from both sequences. Each feature channel was normalized before calculating the corresponding Huber losses. The total loss was

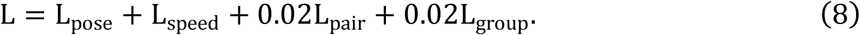

Here, the four terms represent the pose, group-centroid speed, pairwise-feature, and group-configuration reconstruction losses, respectively.

Pose augmentation was applied to 75% of training inputs, while the remaining inputs were left unchanged. During epochs 1–40, zero-mean Gaussian coordinate noise with a standard deviation of 0.004 body lengths was applied together with scale perturbations of ±0.4%. During epochs 41– 100, the noise standard deviation increased to 0.005 body lengths and the scale perturbations to ±0.5%. Training was performed for 100 epochs using the AdamW optimizer (*76*), with a learning rate of 5 × 10⁻⁵, weight decay of 1 × 10⁻⁵, and batch size of 128. A 15-epoch linear warm-up was followed by cosine annealing, and the gradient norm was clipped at 1.0. The random seed was fixed at 42, deterministic training options were enabled, and the checkpoint with the lowest validation loss was selected for downstream analyses.

### Construction of behavioral coordinates

Interpretable target coordinates were constructed from window-level summaries of pair- and group-level features. Directional nose-to-tail distances and facing scores were first averaged across directions within each pair. Pair-level features were then averaged across frames and animal pairs, while group-level features were averaged across frames. Values were clipped at the Control reference 1st and 99th percentiles and standardized using the mean and standard deviation of the clipped Control values.

Block-wise parallel analysis (*77*) determined the number of factors. Principal-axis factor analysis (*78*) with Promax rotation (*79*) yielded three factors based on their loadings. Factor scores were calculated using features with absolute loadings ≥0.40, weighted by their signed loadings divided by the sum of the retained absolute loadings, and standardized within the Control reference.

An MLP with two hidden layers served as a projector from the frozen 64-dimensional DAE embeddings to the three-dimensional behavioral space. Each Control cage was projected using an MLP trained on all other Control cages. Input and target standardization was based on those training cages. The target coordinate system remained fixed across folds. VPA coordinates were obtained by averaging predictions across all Control-trained projectors.

### Behavioral organization index and trajectory analysis

For each cage at each week, the Spacing, Orientation, and Dynamics coordinates were summarized by their medians and interquartile ranges, yielding six behavioral features. A ridge regression model predicted postnatal age from these features using Control data, with nested leave-one-Control-cage-out cross-validation for model evaluation and ridge-penalty selection. The fitted model was applied to the held-out Control cage and all VPA cages, with VPA predictions averaged across outer folds.

The behavioral organization index (BOI) was defined as

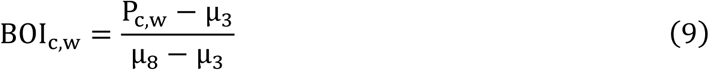

where P_c,w_ is the predicted age for cage *c* at observed week *w*, and μ_3_and μ_8_ are the mean predicted ages across Control cages at 3 and 8 weeks, respectively. This scaling set the mean Control BOI to 0 at 3 weeks and 1 at 8 weeks. Early-period BOI was summarized as the trapezoidal area under the BOI trajectory from 3 to 6 weeks, divided by the three-week interval.

Longitudinal BOI was analyzed using a linear mixed-effects model fitted by restricted maximum likelihood. Fixed effects included condition, categorical week, and their interaction. The model included nested random intercepts for dam, litter, and cage, with multiplicative condition- and week-specific residual standard deviations. Model contrasts were evaluated using dam-clustered CR1 standard errors and Satterthwaite degrees of freedom.

### Motif definition and transition analysis

Motifs were defined in the three-dimensional behavioral coordinate space using a Gaussian mixture model (GMM) fitted exclusively to Control data. Four components were used based on their behavioral distinctiveness and reproducibility across repeated fits and the components were labeled M0–M3 according to their positions along the Spacing, Orientation, and Dynamics axes. The fitted model was applied to all Control and VPA recordings to estimate motif posterior probabilities for each window. Occupancy of each motif was calculated as its mean posterior probability across windows within each cage-week.

For transition analysis, each window was assigned to the motif with the highest posterior probability. Transitions were counted between adjacent 1-s windows within each recording. Windows separated by excluded intervals were not treated as adjacent. Counts were summed across recordings within each cage-week, including self-transitions. The transition probability from motif *a* to motif *b* was calculated as

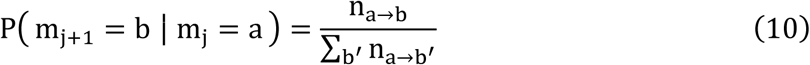

where n_a→b_ is the transition count within a cage-week, and the denominator sums transitions from motif *a* to all destination motifs, including itself. Group-level transition probabilities were calculated separately at each week by averaging cage-level probabilities. Age effects were tested on square-root-transformed joint distributions using within-cage permutations of week.

### Statistical analysis

Statistical analyses were performed using Python and R. Longitudinal BOI mixed-effects models and model contrasts were evaluated in R using dam-clustered CR1 standard errors and Satterthwaite degrees of freedom. Longitudinal home-cage data were analyzed at the cage level, with dam, litter, and cage clustering accounted for in BOI mixed-effects analyses. Mother–pup associations with BOI were assessed at the litter level, with offspring BOI averaged across sibling postweaning cages where applicable. Individual mice and videos were the units of analysis for task-based behavioral assays and tracking-performance comparisons, respectively.

Unless otherwise specified, statistical tests were two-sided, and P < 0.05 was considered statistically significant. Multiple-comparison corrections were applied as specified for each analysis. Benjamini–Hochberg-adjusted P values were reported as q values, with q < 0.05 considered statistically significant. Exact sample sizes, statistical tests, test statistics, and P or q values are provided in the corresponding figure legends.

## Supporting information

Movie S1. Tracking across mouse recording settings

Movie S2. Tracking across additional animal species

Movie S3. Examples of behavioral motifs

## Author contributions

Conceptualization: S.K., A.C.

Software Development and Data management: S.K., Y.J.

Experimental Investigation: S.K., H.L.T., H.C.

Resources: A.C.

Project administration: S.K., A.C.

Supervision: A.C.

Funding acquisition: A.C.

Writing: S.K., Y.J., and A.C.

Visualization: S.K., Y.J.

## Data, code, and materials availability

MovAl source code used in this study (version paper-2026-09-17) is archived at https://doi.org/10.5281/zenodo.22810176. Analysis code for Figs. 2–5 (Behavior_traj, version v1.0.0) is archived at https://doi.org/10.5281/zenodo.22810180. The corresponding GitHub repositories are https://github.com/coldlabkaist/MovAl and https://github.com/coldlabkaist/Behavior_traj. CSV datasets used in the analyses are available at https://drive.google.com/drive/folders/1daqqVoYe7smMXBZlpLwBUThRYb_N6N0e. Original video recordings are available upon request from the corresponding author, Ain Chung. This study did not generate any new materials.

