## Supplementary Materials for "Developmental Dynamics of Naturalistic Social Behavior Revealed by Longitudinal Multi-Animal Tracking"

Sunjin Kim et al.

**This PDF file includes:**

Figs. S1 to S10
Tables S1 to S2
Legends for movies S1 to S4

**Other Supplementary Materials for this manuscript include the following:**

Movies S1 to S4

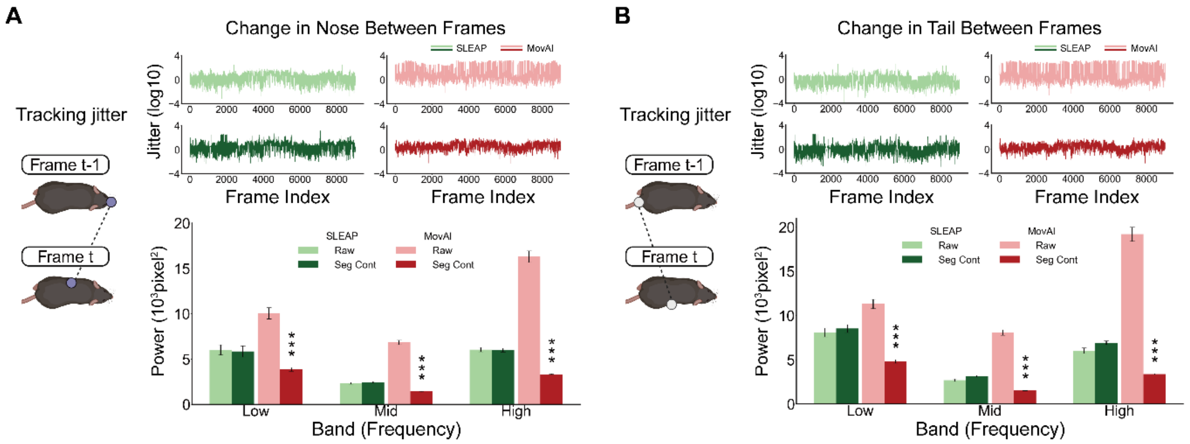


**Fig. S1. Tracking-jitter analysis for nose and tail key points.**

**(A)** Nose tracking jitter. (Top) Representative log10-transformed frame-to-frame displacement time series of the nose key point for SLEAP and MovAl under Raw and Seg-Cont input conditions. (Bottom) Fast Fourier transform–derived jitter power summarized across low-, mid-, and high-frequency bands. **(B)** Tail tracking jitter. (Top) Representative log10-transformed frame-to-frame displacement time series of the tail key point for each method and input condition. (Bottom) Fast Fourier transform–derived jitter power summarized across the three frequency bands. For both panels, bars and error bars indicate the mean ± SEM across 17 videos after averaging the three tracks within each video (two-way repeated-measures ANOVA for each key point and frequency band, followed by two-sided paired t tests with Holm correction across the six pairwise comparisons). Asterisks above Seg-Cont MovAl indicate comparisons with Raw MovAl. Detailed Holm-adjusted P values for all pairwise comparisons are provided in Supplementary Table 2. ***$P < 0.001$. The BioRender credit applies to the mouse illustrations. Created in BioRender. Kaist, C. (2026) https://BioRender.com/19belk0

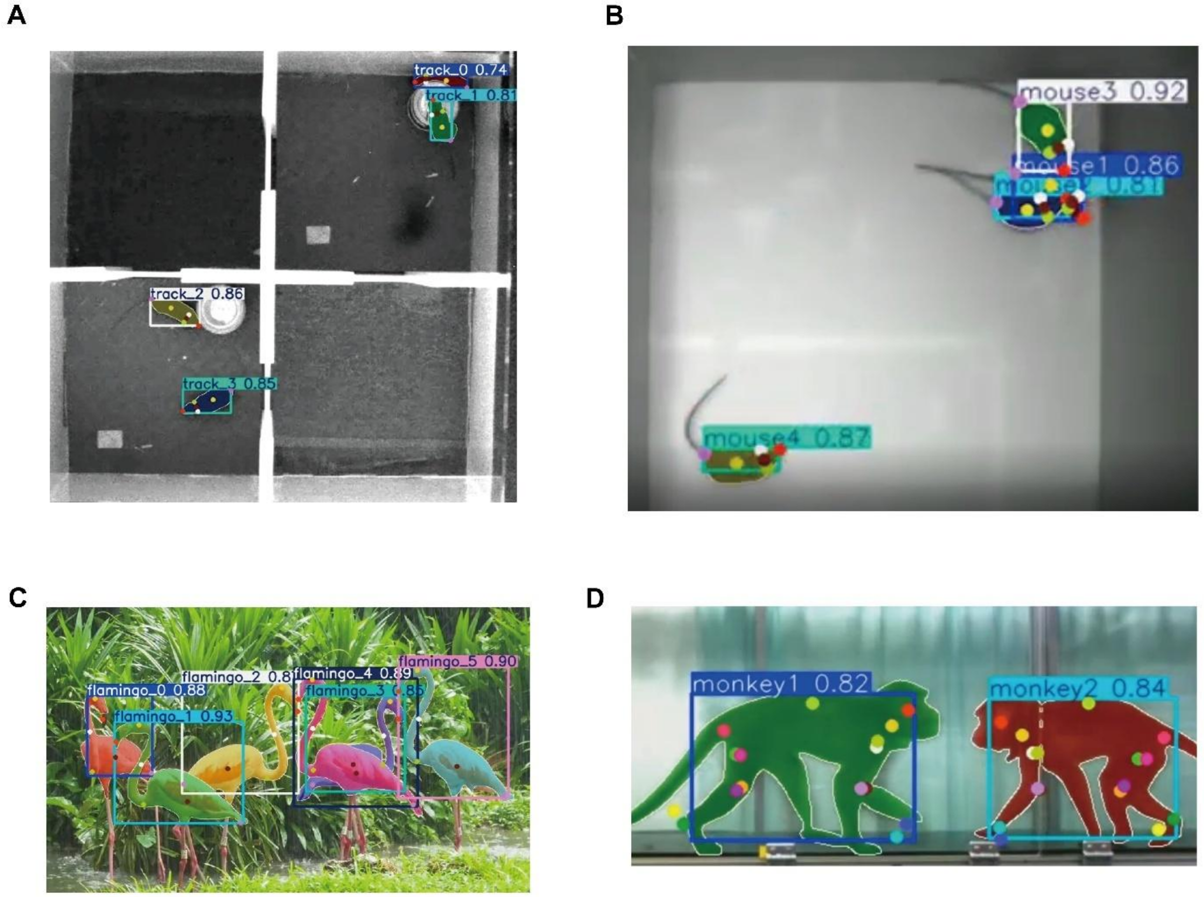


**Fig. S2. Application of MovAl across diverse experimental settings.**

Representative MovAl outputs are shown with segmentation masks, pose keypoints, identity labels, and detection confidence scores. **(A)** Simultaneous tracking of four mice in a chamber divided into two compartments, with two mice housed in each compartment. **(B)** Tracking of four freely moving mice in an open-field arena. **(C)** Six flamingos were recorded from a side view. **(D)** Tracking of two freely interacting monkeys. These examples illustrate the applicability of MovAl across variations in enclosure configuration, animal number, image quality, physical appearance, and species.

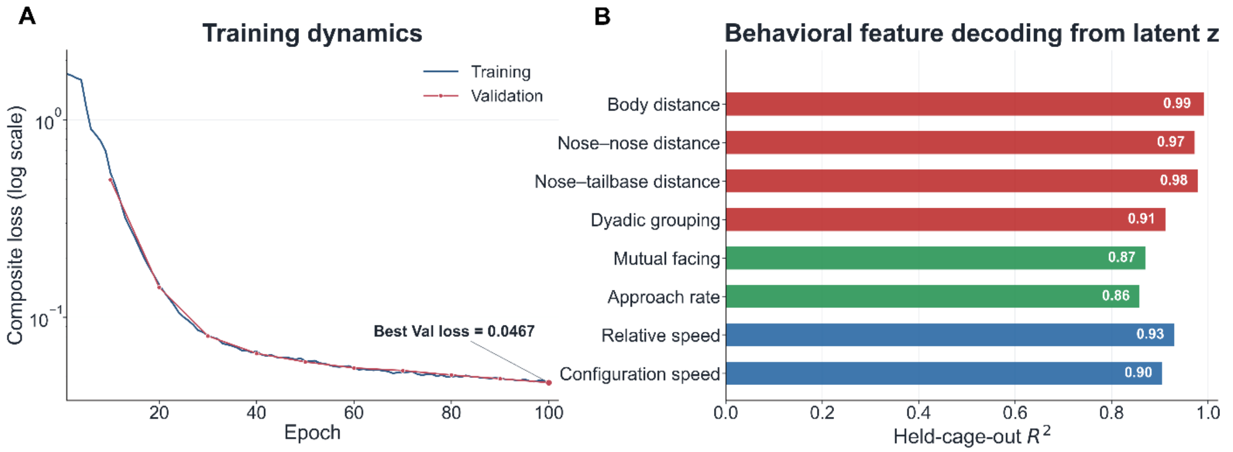


**Fig. S3. DAE training dynamics and validation of the latent representation.**

**(A)** Training and validation composite losses decreased in parallel over 100 epochs, reaching a best validation loss of 0.0467. The close agreement between the two curves indicated stable model convergence without substantial divergence between the training and validation data. Loss is shown on a logarithmic scale. **(B)** Behavioral information retained in the frozen sequence-level latent representation z was evaluated by predicting eight pose-derived features under held-cage-out validation. Spatial-relationship features were decoded with $R^2$ values of 0.91–0.99, orientation- and approach-related features with $R^2$ values of 0.86–0.87, and movement-dynamics features with $R^2$ values of 0.90–0.93. These results demonstrate that the latent representation preserved information about inter-animal configuration, social orientation, and group movement across unseen cages, supporting its subsequent mapping into an interpretable behavioral space. Red, green, and blue bars indicate spatial-relationship, orientation/approach, and movement-dynamics features, respectively.

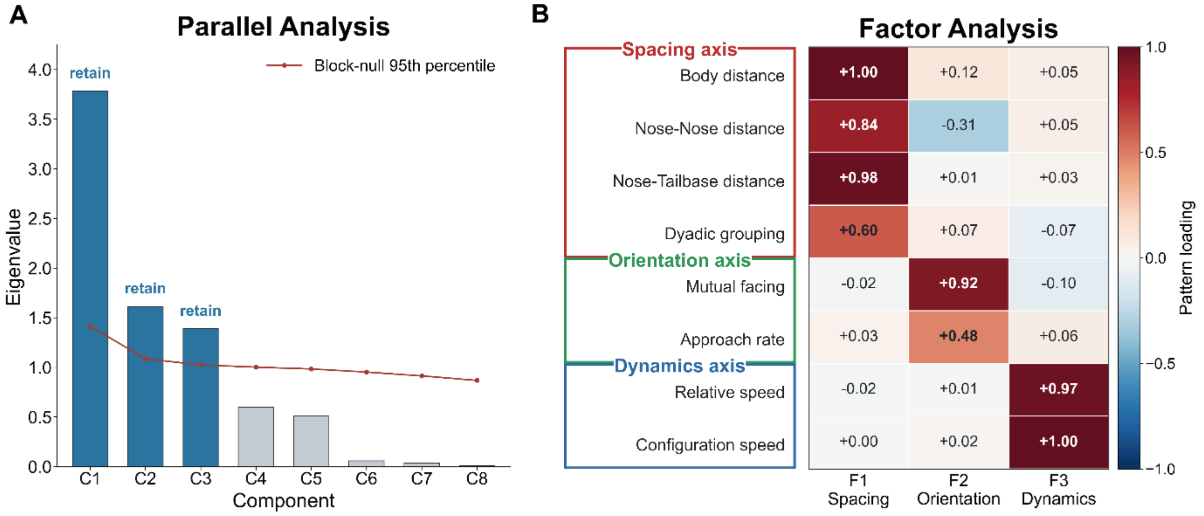


**Fig. S4. Parallel analysis and factor structure of pose-derived behavioral features.**

**(A)** Block-wise parallel analysis used to determine the number of factors to retain. Eigenvalues obtained from the observed behavioral features are shown as bars and were compared with the 95th percentile of the corresponding block-null distribution (red line). Only the first three components exceeded the null threshold, whereas components C4–C8 fell below it. These results supported retention of a three-factor solution describing inter-animal spacing, social orientation, and interaction dynamics. **(B)** Pattern-loading matrix obtained by principal-axis factor analysis with Promax rotation. The eight pose-derived behavioral features were organized into three interpretable factors. F1 showed strong positive loadings for body distance (+1.00), nose–nose distance (+0.84), nose–tail-base distance (+0.98), and dyadic grouping (+0.60) and was therefore interpreted as the Spacing axis. F2 was primarily defined by mutual facing (+0.92) and approach rate (+0.48), representing the Orientation axis. F3 showed strong loadings for relative speed (+0.97) and configuration speed (+1.00) and was interpreted as the Dynamics axis. Most cross-loadings on the other factors were small, indicating that the three factors captured largely distinct aspects of group behavior. Cell values and colors indicate the magnitude and direction of the pattern loadings.

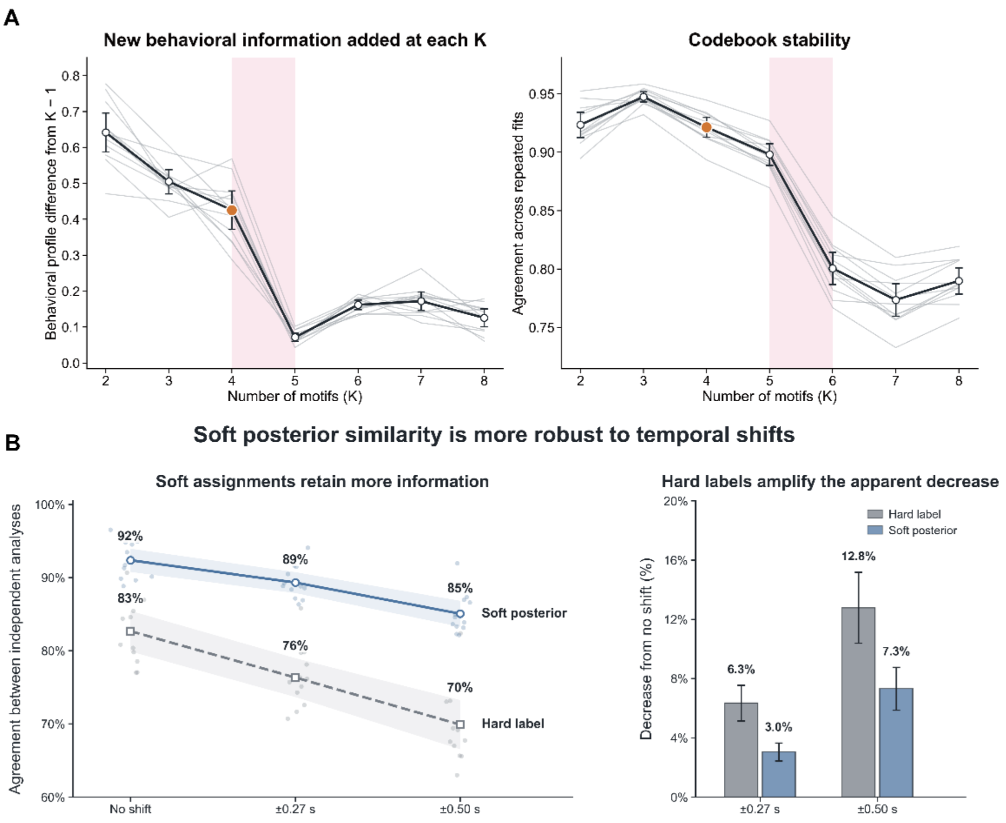


**Fig. S5. Robustness of soft motif assignments and rationale for selecting four motifs.**

**(A)** The number of motifs was evaluated by considering both the behavioral information captured by each additional motif and the reproducibility of the resulting codebook. (Left) The x-axis indicates the number of motifs K, and the y-axis indicates the behavioral-profile difference between solutions with K and K − 1 motifs. A larger value means that increasing K captures an additional, behaviorally distinct pattern. The substantial differences observed through (K=4) indicate that solutions with fewer than four motifs omit meaningful behavioral resolution. In contrast, the marked reduction at (K=5) indicates that motifs added beyond (K=4) contribute little new behavioral information. (Right) The x-axis indicates (K), and the y-axis shows posterior agreement across repeated model fits, with higher values indicating a more reproducible codebook. The four-motif solution maintained high agreement, whereas stability declined as the codebook was divided into more motifs, particularly from (K=5) to (K=6). **(B)** Temporal robustness of motif assignments was assessed by independently repeating the analysis after shifting the behavioral window boundaries. (Left) The x-axis indicates the magnitude of the temporal shift, and the y-axis indicates the agreement between two independently fitted analyses. For soft assignments, agreement was measured as the similarity between posterior probability vectors assigned to corresponding windows. For hard assignments, it was measured as the percentage of windows assigned to the same matched motif. The no-shift condition represents agreement between independent repeated fits and therefore provides a baseline for ordinary model-fitting variability. Soft posterior agreement remained higher than hard-label agreement at both temporal shifts. (Right) The x-axis indicates the magnitude of the temporal shift, and the y-axis shows the percentage-point decrease in agreement relative to the no-shift baseline. Smaller values therefore indicate greater robustness to temporal displacement.

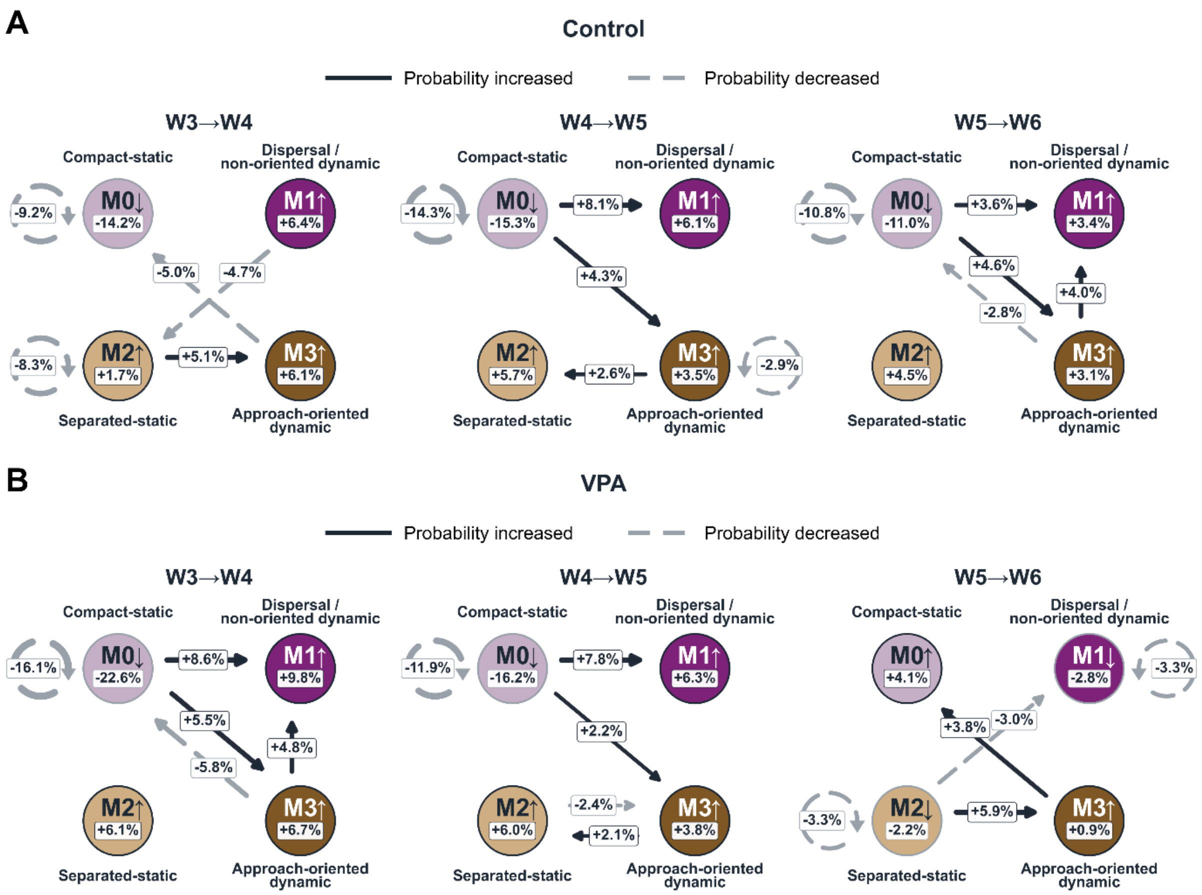


**Fig. S6. Early developmental changes in motif occupancy and transition organization.**

**(A)** Adjacent-week changes across the W3–W4, W4–W5, and W5–W6 intervals in the Control group. **(B)** Corresponding changes in the VPA group. Colored nodes represent the four behavioral motifs, with values inside each node indicating changes in mean soft motif occupancy. Directed edges connect source and destination motifs and display the five largest absolute changes in mean conditional transition probability calculated from hard-label assignments within each interval. Curved arrows that loop back to the same node represent self-transitions, indicating changes in motif recurrence. Node and edge values are expressed in percentage points. Solid dark arrows denote increases, dashed gray arrows denote decreases, and line width increases with the absolute magnitude of the transition change using a common scale across panels.

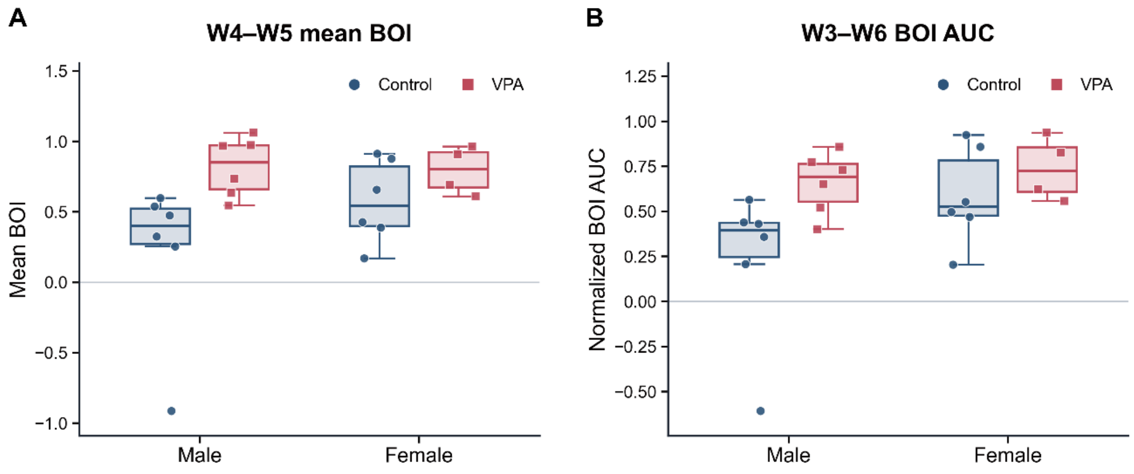


**Fig. S7. Sex-stratified comparisons of early postweaning BOI.**

**(A)** BOI averaged across 4 and 5 weeks within each cage (longitudinal mixed-effects model; *F*-tests with approximate Satterthwaite degrees of freedom and Benjamini–Hochberg correction across the two endpoints for each effect; condition: $F(1, 27.73) = 8.27$, $q = 0.0107$; sex: $F(1, 25.33) = 1.55$, $q = 0.2247$; condition × sex: $F(1, 25.33) = 1.48$, $q = 0.2397$). **(B)** Normalized BOI AUC from 3 to 6 weeks (same model and correction; condition: $F(1, 24.32) = 7.65$, $q = 0.0107$; sex: F(1, 31.20) = 4.82, $q = 0.0714$; condition × sex: $F(1, 31.20) = 1.44$, $q = 0.2397$). Points represent individual cages; boxes indicate the median and interquartile range, and whiskers extend to the most extreme observations within 1.5 times the interquartile range (Control: 6 male and 6 female cages; VPA: 6 male and 4 female cages). The model accounted for dam/litter/cage clustering and condition- and week-dependent residual variances; main effects were equally averaged over the levels of the other factor.

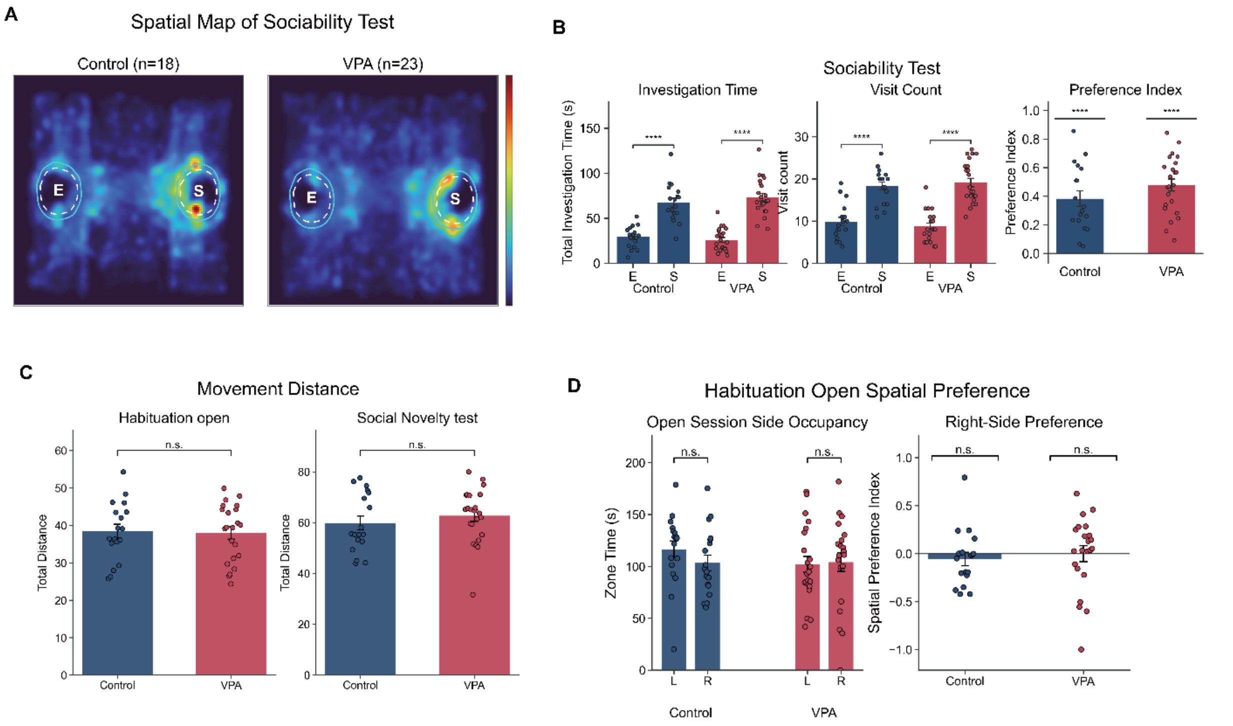


**Fig. S8. Analysis of three-chamber sociability, movement, and spatial-bias.**

**(A)** Group-averaged spatial density maps of nose positions during the sociability phase of the three-chamber test at PND 52. Dashed circles indicate the investigation regions surrounding the empty (E) and social-stimulus (S) cups, and colors from blue to red indicate increasing nose-position density (Control, *n*=18; VPA, *n*=23). **(B)** Sociability quantified by investigation time, visit count, and sociability preference index. Investigation was defined as nose entry into the cup-centered regions of interest. Each point represents one mouse, and bars and error bars indicate the mean ± SEM. (Left) Total investigation time in the empty- and social-cup regions (two-sided paired t-tests: Control, *t*(17)=6.62, *P*<0.0001; VPA, *t*(22)=9.63, *P*<0.0001). (Middle) Number of visits to the empty- and social-cup regions (two-sided paired t-tests: Control, *t*(17)=6.81, *P*<0.0001; VPA, *t*(22)=8.04, *P*<0.0001). (Right) Sociability preference index, calculated as ((S-E)/(S+E)), where (S) and (E) denote investigation times in the social- and empty-cup regions, respectively (two-sided one-sample t-tests against zero: Control, *t*(17)=7.06, *P*<0.0001; VPA, *t*(22)=11.37, *P*<0.0001). **(C)** Total movement distance calculated from body-center trajectories during the habituation and social-novelty phases and expressed in arena-normalized units. Each point represents one mouse, and bars and error bars indicate the mean ± SEM (two-sided Welch's t-tests: habituation, Control, *n*=18, VPA, *n*=22, *t*(36.03)=0.18, *P*=0.861; social-novelty phase, Control, *n*=18, VPA, *n*=23, *t*(35.28)=-0.82, *P*=0.420). **(D)** Spatial side preference during habituation. (Left) Time spent in the left and right compartments (two-sided paired t-tests: Control, *t*(17)=0.90, *P*=0.382; VPA, *t*(21)=-0.14, *P*=0.888). (Right) Right-side preference index, calculated as ((R-L)/(R+L)), where (R) and (L) denote time spent in the right and left compartments, respectively (two-sided one-sample t-tests against zero: Control, *t*(17)=-0.81, *P*=0.430; VPA, *t*(21)=-0.01, *P*=0.990). ****P<0.0001; ns, not significant.

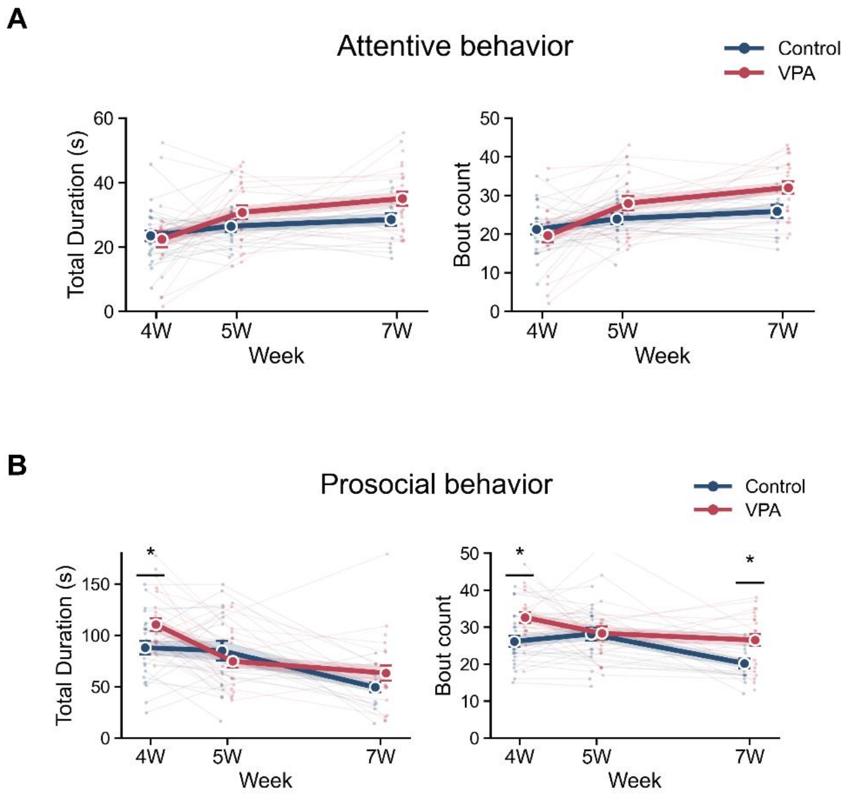


**Fig. S9. Attentive and prosocial behavior of reciprocal social interaction.**

**(A)** Attentive behavior, comprising approach, facing, and following behaviors directed toward a freely moving unfamiliar mouse. (Left) Total duration of attentive behavior (two-sided Welch's t-tests with Benjamini–Hochberg correction across the six comparisons: 4 weeks, $t(38.60)=0.37$, $q=0.712$; 5 weeks, $t(36.69)=-1.54$, $q=0.200$; 7 weeks, $t(35.44)=-2.24$, $q=0.094$). (Right) Attentive-behavior bout count (two-sided Welch's t-tests with Benjamini–Hochberg correction across the six comparisons: 4 weeks, $t(41.00)=0.74$, $q=0.555$; 5 weeks, $t(35.74)=-1.74$, $q=0.182$; 7 weeks, $t(33.84)=-2.74$, $q=0.0581$). **(B)** Prosocial behavior, comprising nose-to-head, nose-to-body, nose-to-anogenital, and mounting behaviors. (Left) Cumulative duration of prosocial behavior (two-sided Welch's t-tests with Benjamini–Hochberg correction across the six comparisons: 4 weeks, $t(44.81)=-2.48$, $q=0.0339$; 5 weeks, $t(29.75)=0.90$, $q=0.451$; 7 weeks, $t(34.57)=-1.56$, $q=0.191$). (Right) Prosocial-behavior bout count (two-sided Welch's t-tests with Benjamini–Hochberg correction across the six comparisons: 4 weeks, $t(44.98)=-3.12$, $q=0.0131$; 5 weeks, $t(36.93)=-0.09$, $q=0.931$; 7 weeks, $t(35.99)=-3.04$, $q=0.0131$). Light lines represent individual mice, and dark lines and error bars indicate the mean ± SEM (Control/VPA: 4 weeks, $n=24/23$; 5 weeks, $n=18/21$; 7 weeks, $n=15/23$). $*q<0.05$.

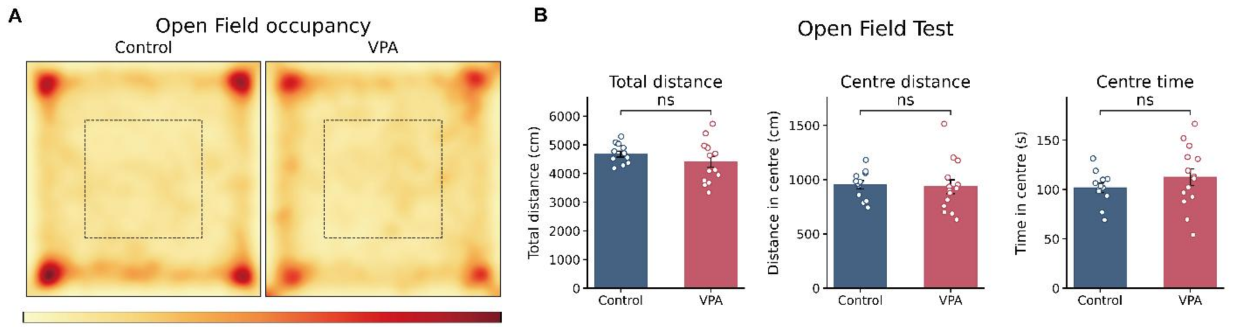


**Fig. S10. Results of open-field test in Control and VPA-exposed mice.**

**(A)** Group-averaged spatial occupancy maps during the open-field test performed at PND 38. The dashed square indicates the center zone, defined as the inner 50% of each spatial axis. Occupancy maps were normalized within each recording before group averaging so that each mouse contributed equally. Control and VPA-exposed mice showed broadly similar spatial distributions, with greater occupancy near the arena periphery and corners. **(B)** Total distance traveled, distance traveled within the center zone, and time spent in the center zone. Each point represents one mouse, and bars and error bars indicate the mean ± SEM (Control, $n$ = 12; VPA, $n$ = 14). (Left) Total distance traveled (two-sided Welch's t-test, $t(19.32) = -1.18$, $P = 0.252$). (Middle) Distance traveled within the center zone (two-sided Welch's t-test, $t(20.83) = -0.22$, $P = 0.828$). (Right) Time spent in the center zone (two-sided Welch's t-test, $t(20.43) = 1.10$, $P = 0.285$). ns, not significant.

**(a) Frequency of tracking miss error in SLEAP, DLC and MovAl**

| Key Point | Raw SLEAP (%) | Seg-Cont SLEAP (%) | Raw DLC (%) | Seg-Cont DLC (%) | Raw MovAl (%) | Seg-Cont MovAl (%) |
|---|---|---|---|---|---|---|
| Nose | 9.63 ± 2.28 | 7.63 ± 1.59 | 11.39 ± 1.96 | 14.14 ± 2.75 | **0.56 ± 0.20** | **0.78 ± 0.23** |
| Ear_L | 8.12 ± 2.01 | 9.43 ± 1.87 | 7.09 ± 1.73 | 10.41 ± 2.85 | **0.56 ± 0.20** | **0.78 ± 0.23** |
| Ear_R | 8.06 ± 2.00 | 9.34 ± 2.11 | 6.75 ± 1.54 | 9.82 ± 2.44 | **0.56 ± 0.20** | **0.78 ± 0.23** |
| Body_C | 6.85 ± 1.82 | 2.29 ± 1.04 | 5.82 ± 1.47 | 7.85 ± 2.36 | **0.56 ± 0.20** | **0.78 ± 0.23** |
| Tail | 14.79 ± 3.82 | 6.02 ± 1.16 | 10.11 ± 1.70 | 20.41 ± 3.33 | **0.56 ± 0.20** | **0.78 ± 0.23** |

**(b) Statistical summary of tracking miss error frequency across tracking methods.**

| Key Point | Group1 | Group2 | Holm-adjusted P |
|---|---|---|---|
| Nose | Raw SLEAP | Seg-Cont SLEAP | 0.654 |
| | | Raw DLC | 0.654 |
| | | Seg-Cont DLC | 0.310 |
| | | **Raw MovAl** | **0.006** |
| | | **Seg-Cont MovAl** | **0.006** |
| | Seg-Cont SLEAP | Raw DLC | 0.151 |
| | | Seg-Cont DLC | 0.094 |
| | | **Raw MovAl** | **0.003** |
| | | **Seg-Cont MovAl** | **0.003** |

| | | | |
|---|---|---|---|
| | Raw DLC | Seg-Cont DLC | 0.310 |
| | | **Raw MovAl** | **< 0.001** |
| | | **Seg-Cont MovAl** | **< 0.001** |
| | Seg-Cont DLC | **Raw MovAl** | **0.001** |
| | | **Seg-Cont MovAl** | **0.001** |
| | Raw MovAl | Seg-Cont MovAl | 0.121 |
| Ear_L | Raw SLEAP | Seg-Cont SLEAP | 1.000 |
| | | Raw DLC | 1.000 |
| | | Seg-Cont DLC | 1.000 |
| | | **Raw MovAl** | **0.013** |
| | | **Seg-Cont MovAl** | **0.014** |
| | Seg-Cont SLEAP | Raw DLC | 0.665 |
| | | Seg-Cont DLC | 1.000 |
| | | **Raw MovAl** | **0.002** |
| | | **Seg-Cont MovAl** | **0.002** |
| | Raw DLC | Seg-Cont DLC | 0.665 |
| | | **Raw MovAl** | **0.014** |
| | | **Seg-Cont MovAl** | **0.014** |
| | Seg-Cont DLC | **Raw MovAl** | **0.022** |
| | | **Seg-Cont MovAl** | **0.022** |
| | Raw MovAl | Seg-Cont MovAl | 0.141 |
| Ear_R | Raw SLEAP | Seg-Cont SLEAP | 1.000 |
| | | Raw DLC | 1.000 |
| | | Seg-Cont DLC | 1.000 |
| | | **Raw MovAl** | **0.012** |

| | | | |
|---|---|---|---|
| | | **Seg-Cont MovAl** | **0.012** |
| | Seg-Cont SLEAP | Raw DLC | 0.825 |
| | | Seg-Cont DLC | 1.000 |
| | | **Raw MovAl** | **0.007** |
| | | **Seg-Cont MovAl** | **0.007** |
| | Raw DLC | Seg-Cont DLC | 0.656 |
| | | **Raw MovAl** | **0.008** |
| | | **Seg-Cont MovAl** | **0.008** |
| | Seg-Cont DLC | **Raw MovAl** | **0.012** |
| | | **Seg-Cont MovAl** | **0.012** |
| | Raw MovAl | Seg-Cont MovAl | 0.141 |
| Body_C | Raw SLEAP | Seg-Cont SLEAP | 0.298 |
| | | Raw DLC | 1.000 |
| | | Seg-Cont DLC | 1.000 |
| | | **Raw MovAl** | **0.028** |
| | | **Seg-Cont MovAl** | **0.029** |
| | Seg-Cont SLEAP | Raw DLC | 0.301 |
| | | Seg-Cont DLC | 0.298 |
| | | **Raw MovAl** | 0.541 |
| | | **Seg-Cont MovAl** | 0.606 |
| | Raw DLC | Seg-Cont DLC | 0.884 |
| | | **Raw MovAl** | **0.026** |
| | | **Seg-Cont MovAl** | **0.028** |
| | Seg-Cont DLC | **Raw MovAl** | **0.055** |
| | | **Seg-Cont MovAl** | **0.055** |

| | | | |
|---|---|---|---|
| | Raw MovAl | Seg-Cont MovAl | 0.181 |
| Tail | Raw SLEAP | Seg-Cont SLEAP | 0.131 |
| | | Raw DLC | 0.527 |
| | | Seg-Cont DLC | 0.527 |
| | | **Raw MovAl** | **0.012** |
| | | **Seg-Cont MovAl** | **0.012** |
| | Seg-Cont SLEAP | Raw DLC | 0.124 |
| | | Seg-Cont DLC | **0.002** |
| | | **Raw MovAl** | **0.001** |
| | | **Seg-Cont MovAl** | **0.002** |
| | Raw DLC | Seg-Cont DLC | **0.004** |
| | | **Raw MovAl** | **< 0.001** |
| | | **Seg-Cont MovAl** | **< 0.001** |
| | Seg-Cont DLC | **Raw MovAl** | **< 0.001** |
| | | **Seg-Cont MovAl** | **< 0.001** |
| | Raw MovAl | Seg-Cont MovAl | 0.101 |

**Table S1. Tracking miss error frequency across tracking methods**

Tracking miss errors across key points, with a statistical summary of tracking miss error frequency across tracking methods; statistical significance was assessed using a two-way repeated-measures ANOVA followed by Holm-adjusted post-hoc comparisons.

| Group | | p-value | | |
|---|---|---|---|---|
| **Group1** | **Group2** | **Body_C** | **Nose** | **Tail** |
| (a) Low Bandwidth (0.00–0.05) | | | | |
| Raw SLEAP | Seg-Cont SLEAP | 0.113 | 0.692 | 0.239 |
| Raw SLEAP | Raw MovAl | < 0.001 | < 0.001 | < 0.001 |
| Raw SLEAP | Seg-Cont MovAl | < 0.001 | 0.012 | < 0.001 |
| Seg-Cont SLEAP | Raw MovAl | 0.013 | < 0.001 | < 0.001 |
| Seg-Cont SLEAP | Seg-Cont MovAl | < 0.001 | 0.003 | < 0.001 |
| Raw MovAl | Seg-Cont MovAl | < 0.001 | < 0.001 | < 0.001 |
| (b) Mid Bandwidth (0.05–0.15) | | | | |
| Raw SLEAP | Seg-Cont SLEAP | 0.092 | 0.221 | 0.036 |
| Raw SLEAP | Raw MovAl | < 0.001 | < 0.001 | < 0.001 |
| Raw SLEAP | Seg-Cont MovAl | < 0.001 | < 0.001 | < 0.001 |
| Seg-Cont SLEAP | Raw MovAl | < 0.001 | < 0.001 | < 0.001 |
| Seg-Cont SLEAP | Seg-Cont MovAl | < 0.001 | < 0.001 | < 0.001 |
| Raw MovAl | Seg-Cont MovAl | < 0.001 | < 0.001 | < 0.001 |
| (c) High Bandwidth (0.15–0.50) | | | | |
| Raw SLEAP | Seg-Cont SLEAP | 0.782 | 0.797 | 0.098 |
| Raw SLEAP | Raw MovAl | < 0.001 | < 0.001 | < 0.001 |
| Raw SLEAP | Seg-Cont MovAl | < 0.001 | < 0.001 | < 0.001 |
| Seg-Cont SLEAP | Raw MovAl | < 0.001 | < 0.001 | < 0.001 |
| Seg-Cont SLEAP | Seg-Cont MovAl | < 0.001 | < 0.001 | < 0.001 |
| Raw MovAl | Seg-Cont MovAl | < 0.001 | < 0.001 | < 0.001 |

**Table S2. Statistical summary of tracking jitter spectral power**

Comparison of tracking-jitter spectral power across tracking methods and input conditions. Statistical significance was assessed using a two-way repeated-measures ANOVA for each key point and frequency band, followed by two-sided paired t-tests with Holm correction across the six pairwise comparisons within each band. Analyses used 17 videos, with the three tracks averaged within each video. Values in the table are Holm-adjusted P values.

**Movie S1. Examples of MovAl tracking across mouse recording settings.**

Representative tracking outputs for four mice in a divided chamber, an open-field arena, a home cage, and an arena with reflective walls. Overlays show segmentation masks, keypoints, identity labels, and confidence scores. These clips provide qualitative examples.

**Movie S2. Examples of MovAl tracking across additional animal species.**

Representative tracking outputs for two monkeys, six flamingos, and five puppies. Overlays show segmentation masks and pose keypoints, with identity labels and confidence scores where displayed. These clips illustrate potential applicability across species.

**Movie S3. Examples of the four behavioral motifs identified in the behavioral space.**

Example behavioral sequences illustrate the four motifs identified by fitting a Gaussian mixture model to Control behavioral windows (Fig. 4C). Columns show M0 (compact-static), M1 (dispersal/non-oriented dynamic), M2 (separated-static), and M3 (approach-oriented dynamic). The upper row shows video recordings with pose overlays, and the lower row shows the corresponding pose representations. These examples illustrate differences in inter-animal spacing, orientation, and movement across motifs.

**Movie S4. Examples of MovAl tracking in mother–pup recordings at PNDs 10, 15, and 20.**

Representative home-cage recordings showing simultaneous tracking of a mother and pups at postnatal days (PNDs) 10, 15, and 20. Segmentation masks and pose keypoints are overlaid on the recordings to illustrate the tracking underlying the mother–pup analyses (Fig. 5E).
